# Quantitative proximity proteomics resolves the epithelial apical-lateral border and uncovers a vertebrate marginal zone defined by the polarity protein Pals1

**DOI:** 10.1101/710202

**Authors:** Benedict Tan, Suat Peng, Sara Sandin, Jayantha Gunaratne, Walter Hunziker, Alexander Ludwig

**Affiliations:** Epithelial Cell Biology Laboratory, Institute of Molecular and Cell Biology, Agency for Science, Technology and Research (A*STAR), Singapore; Quantitative Proteomics Group, Institute of Molecular and Cell Biology, Agency for Science, Technology and Research (A*STAR), Singapore; Department of Physiology, Yong Loo Lin School of Medicine, National University of Singapore; Department of Anatomy, Yong Loo Lin School of Medicine, National University of Singapore; School of Biological Sciences, Nanyang Technological University, Singapore; NTU Institute of Structural Biology, Nanyang Technological University, Singapore

## Abstract

Epithelial apico-basal polarity is established through the asymmetric cortical distribution of the Par, Crumbs and Scribble polarity modules. Apical (Par and Crumbs) and basolateral (Scribble) polarity modules overlap at the apical-lateral border, which, in mammals, is defined by the apical junctional complex (AJC). The AJC is composed of tight junctions (TJ) and adherens junctions (AJ) and plays fundamental roles in epithelial morphogenesis and plasticity. However, the molecular composition and precise sub-junctional organization of the AJC and its associated polarity regulators are still not well defined. Here we used the peroxidase APEX2 for quantitative proximity proteomics (QPP) and electron microscopy (EM) imaging to generate a nanometer-scale spatio-molecular map of the apical-lateral border in fully polarized MDCK-II cells. Using Par3 and Pals1 as surrogates for QPP we present a spatially resolved network of ∼800 junction-associated proteins. The network dissects TJ and AJ components and provides strong evidence that TJ are composed of distinct apical and basal subdomains. Moreover, we find that Pals1 and its binding partners PatJ, Lin7c and Crumbs3 define a hitherto unidentified membrane compartment apical of TJ, which we coin the vertebrate marginal zone (VMZ). The VMZ is physically associated with HOMER scaffolding proteins, regulators of apical exocytosis, and membrane-proximal HIPPO pathway proteins. Taken together our work defines the spatial and molecular organization of the apical-lateral border in fully polarized mammalian epithelial cells, reveals an intriguing molecular and spatial conservation of invertebrate and vertebrate cell polarity protein domains, and provides a comprehensive resource of potentially novel regulators of cell polarity and the mammalian AJC.

## Introduction

Cell polarity is a fundamental process critical for development and for many cell and tissue functions. Epithelial polarity is governed by the evolutionarily conserved polarity modules Par, Scribble and Crumbs, which define distinct apical and basal compartments. While Par3, Par6, atypical PKC (aPKC), Crumbs (Crb), Pals1 (protein associated with Lin7, also known as MPP5) and PatJ (Pals1-associated tight junction protein) constitute the apical polarity machinery, the Scribble module (Scribble (SCRIB), Disks Large (DLG) and Lethal Giant Larvae (LGL)) operates at the lateral membrane [1, 2]. This asymmetric cortical distribution of the polarity proteins is paramount for polarity development, as well as for the formation and correct positioning of epithelial cell junctions [3–8]. In mammalian epithelia, apical and basal polarity modules overlap at the apical junctional complex (AJC), a multi-functional and highly dynamic membrane compartment that demarcates the apical-lateral border. The AJC is composed of apical tight junctions (TJ) and more basally located adherens junctions (AJ). While TJ act as a diffusion barrier and gate to regulate apico-basal polarity and transepithelial permeability, AJ are cell-cell adhesion plaques that control tissue mechanics through the junction-associated actomyosin network [9, 10]. In addition, the AJC acts as a major signaling and trafficking hub to control epithelial morphogenesis and plasticity. How the AJC integrates such a range of biological processes is poorly understood, in part because the full complement of its components as well as its sub-junctional molecular organization are not well defined. In addition, it is unclear how apical and baso-lateral polarity modules are organized at the apical-lateral border, and how their cortical organization relates to the AJC.

In addressing this question, it is informative to consider the organization of the polarity proteins in invertebrate epithelia, which possess a markedly different apico-basal configuration of cell-cell junctions. In particular, invertebrate epithelia (which are best characterized in *Drosophila*) possess a specialized membrane compartment called the marginal zone (MZ, or subapical region). The MZ is located at the apical-most position of the lateral membrane and is defined by a protein complex composed of the fly homologues of Crb, Pals1 (Stardust) and PatJ (Discs Lost) [11]. Par6 and aPKC are also associated with the MZ as well as with the free apical membrane. By contrast, Bazooka (the fly homologue of Par3) is excluded from the MZ and localizes basally of Crb and Stardust, at the zonula adherens (ZA) [12–14]. Thus, in flies, Par3 and components of the MZ define distinct junctional compartments. This is in apparent contrast to vertebrate epithelia, in which both the Crumbs (Crb3, Pals1, PatJ, Lin7) [6, 7, 15–18] and the Par modules (Par3, Par6, aPKC) [8, 19] are thought to localize to TJ. This has led to the hypothesis that the TJ represents a hybrid compartment that has evolved from components of the MZ, the ZA, and septate junctions [10, 11]. However, the precise sub-junctional organization of the mammalian apical polarity proteins has not been studied with sufficient spatial resolution and molecular detail to support such a model. Therefore, whether vertebrates possess a membrane domain related to the MZ, as suggested previously [20], remains unclear.

Proximity proteomics is a powerful approach to generate proteomes of subcellular organelles and to dissect protein-protein interaction (PPI) networks *in situ* [21, 22]. The technique, therefore, appears ideally suited to resolve the molecular and spatial organization of the apical-lateral border. Indeed, proximity proteomes for the TJ proteins ZO-1, occludin, claudin-4 [23, 24] and the AJ protein E-cadherin [25, 26] have been generated using the promiscuous biotin ligase BirA* (BioID). However, all of these proteomes are derived from epithelial cells grown on plastic, and hence provide only limited information on the composition of TJ and AJ in fully polarized epithelia. Moreover, all TJ and AJ proteomes determined to date suffer from significant “cross-contamination”, i.e. TJ proteomes contain bona-fide AJ components, and *vice versa*. This is likely to be due to the extensive labeling times required for BirA* (up to 24 hours), which, given the dynamic nature of the AJC, can lead to off-target labeling in neighboring compartments. As a consequence, the spatial organization of the reported TJ- and AJ-associated proteomes remains elusive. Similarly, although a number of approaches have been employed to identify novel components of the polarity network [27, 28] including pull-down assays [29], yeast two hybrid screens [30], and affinity purification followed by mass spectrometry (AP-MS) [31], such approaches are inherently blind to the spatial organization of the identified PPIs. In addition, low-affinity, short-lived or low-abundant interactions often remain unidentified. Hence, dissecting the molecular architecture of the AJC and its associated polarity machinery requires tools that a) capture the junction-associated proteome *in situ* with high temporal precision, and b) resolve its components along the apico-basal axis at the nanometer level.

Originally designed as a genetic reporter for EM [32], the ascorbate peroxidase APEX2 has become a popular tool for proximity labeling with biotin. APEX2 offers several advantages over BirA*, including rapid labeling kinetics (<1 min), a short half-life (<1 ms) of the biotin phenoxyl radical that is generated during the labeling reaction, and a small biotinylation radius (∼20 nm). APEX2, therefore, reports on the localization of a protein of interest with low nanometer precision, and acquires instant snapshots of the protein’s molecular environment. Here we have used APEX2 for EM imaging and quantitative proximity proteomics (QPP) to dissect the molecular and spatial organization of the AJC and the junction-associated polarity network in fully polarized, filter-grown MDCK-II cells. Our work resolves the molecular architecture of the AJC and uncovers a vertebrate marginal zone apical of TJ that is defined by the polarity proteins Pals1, PatJ, Lin7c, and Crb3.

## Results

### Mammalian Pals1 localizes to a marginal zone apical of tight junctions

We set out to localize Par3 and Pals1 by EM using clonal MDCK-II cell lines expressing APEX2-EGFP fusion proteins (henceforth referred to as Par3-A2E, and Pals1-A2E, respectively). Par3-A2E expressed at levels comparable to endogenous Par3 colocalized with the TJ marker ZO-1 by confocal microscopy (Figure 1A and 1C), as expected. By contrast, when Pals1-A2E was expressed at levels comparable to endogenous Pals1 (Figure 1B, clone #1), we were unable to detect the fusion protein by confocal microscopy. In addition, Pals1-A2E expressed at this level did not produce sufficient contrast in EM, nor did we find any evidence for efficient proximity biotinylation (data not shown). To meet those key requirements, we generated a Pals1-A2E cell line expressing the fusion protein at about five times the level of endogenous Pals1 (Figure 1B, clone #2). Pals1-A2E expressed at this level colocalized with ZO-1 by confocal microscopy (Figure 1D). Weakly stained puncta of Pals1-A2E were also observed in the apical membrane. In addition, both Par-A2E and Pals1-A2E colocalized with PatJ and were found apical of β-catenin (Figure S1A and S1B). The localization of Par3-A2E and Pals1-A2E was identical to that of the respective endogenous proteins in MDCK-II cells cultured under identical conditions (Figure S1C). Hence, both Par3 and Pals1 localize to TJ as determined by diffraction-limited microscopy.

**Figure 1:**
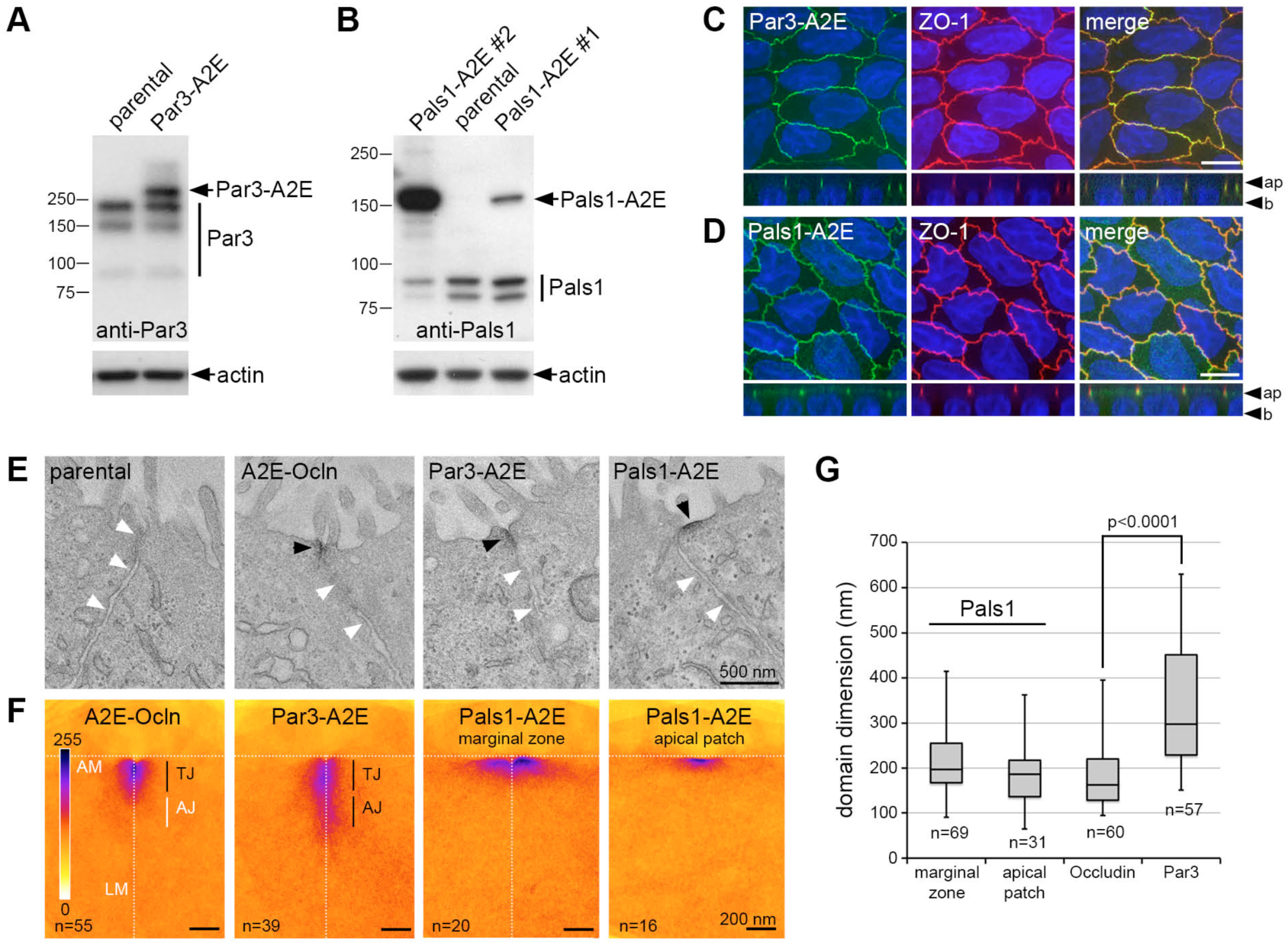
Pals1 is localized to a distinct domain at the margins of the apical membrane. **A and B.** Immunoblots of stable MDCK-II cell lines expressing Par3 (A) or Pals1 (B) APEX2 EGFP fusion proteins (Par3-A2E and Pals1-A2E). Membranes were probed with the indicated antibodies. Note that endogenous Pals1 levels are reduced in Pals1-A2E clone #1. **C and D.** Confocal micrographs of stable Par3-A2E and Pals1-A2E (#2) MDCK-II cells grown on filter membranes for 10 days. Cells were stained with the indicated antibodies and DAPI. Maximum intensity projections are shown. Ap: apical; b: basal. Scale bars are 10 µm. **E.** Representative electron micrographs of untransfected parental MDCK-II cells and MDCK-II cells stably expressing A2E-Occludin (A2E-Ocln), Par3-A2E or Pals1-A2E. Cells were grown on filter membranes and processed for APEX2-EM 10 days post-confluence. **F.** Heat maps showing the localization patterns of APEX2 fusion proteins. Individual EM micrographs were superimposed, aligned and pseudo-coloured. **G.** Domain dimensions of Par3-A2E, Pals1-A2E and A2E-Ocln based on APEX2 labeling. Black arrowheads point to APEX2 staining, white arrowheads indicate the lateral membrane. AM: apical membrane; LM: lateral membrane. Scale bars are indicated. See also Figures S1, S2, and S3.

We then examined the localization of Par3-A2E and Pals1-A2E by EM (Figure 1E and Figure S2). As a spatial landmark, we generated a cell line expressing the integral TJ protein occludin fused to APEX2-EGFP (A2E-Ocln) (Figure S1D and S1E). Electron micrographs from the same specimen were overlaid, aligned and averaged to generate a semi-quantitative heat map of APEX2-mediated EM contrast for each fusion protein (Figure 1F). In addition, the domain dimensions were determined by measuring the extent of labeling along the membrane (Figure 1G). This comprehensive EM analysis revealed distinct localizations and membrane distribution profiles for all three proteins analyzed. A2E-Ocln was localised to the very apical tip of the lateral membrane, extending 183 ± 11 nm (mean ± s.e.m.; n=60) into the lateral membrane. This is consistent with the dimensions of TJ in MDCK cells and with the localization of occludin to TJ strands determined by immuno-gold EM [33, 34]. Par3-A2E localization was often indistinguishable from that of A2E-Ocln, as expected [8, 19]. However, we also observed junctions in which the distribution of Par3-A2E was markedly broader. On average, the Par3 domain extended 340 ± 20 nm (mean ± s.e.m.; n=57) into the lateral membrane, indicating that at least in MDCK-II cells Par3 can localize both to TJ and AJ. In striking contrast to Par3-A2E and A2E-Ocln, Pals1-A2E was entirely absent from apical cell junctions, and instead labeled a distinct compartment at the very margin of the apical membrane. This apical marginal band was 213 ± 11 nm wide (mean ± s.e.m,; n=69) and positioned immediately apical of TJ. In addition, Pals1-A2E labeled discrete patches in the free apical membrane of similar dimensions (183 ± 13 nm (mean ± s.e.m.; n=31)). The localization of all APEX2 fusion proteins was confirmed in EM samples lacking any counterstain (Figure S3). Thus, the observed EM contrast can be attributed solely to the deposition of an electrondense osmiophilic stain that is generated through the local enzymatic activity of the respective APEX2 fusion protein. Taken together our EM analyses provided strong evidence that Par3 and Pals1 occupy distinct membrane domains at the apical-lateral border. While Par3 localised along the length of the AJC, Pals1 was restricted to a hitherto unidentified domain apical of TJ. We henceforth refer to this domain as the ‘vertebrate marginal zone’ (VMZ).

### The VMZ is made of a stable protein complex composed of Pals1, PatJ, Lin7 and Crb3, and its formation requires Par3

Several studies in fly and mammalian epithelia have established physical and functional interactions between the Par and Crb modules [12, 35–37]. Our EM data, however, suggested that in fully polarized MDCK-II cells the Par and Crb modules assemble into separate protein complexes. To test this directly, we immunoprecipitated Pals1-A2E and Par3-A2E from cell lysates of filter-grown and fully polarized MDCK-II cells using anti-GFP antibodies (Figure 2A). We found that aPKC co-immunoprecipitated specifically with Par3-A2E, while PatJ, Lin7c, and Crb3 were all specifically associated with Pals1-A2E. Consistent with previous work [18, 37, 38], Par6 was also specifically recovered in the Pals1-A2E immunoprecipitate, albeit with lower efficiency. Occludin, ZO-2, and actin were not co-immunoprecipitated, confirming the specific nature of the identified interactions. We concluded that Par3 and Pals1 form distinct protein assemblies in fully polarized MDCK-II cells. In addition, the VMZ is likely composed of a stable protein complex containing Pals1, PatJ, Lin7c and Crb3. Next we explored whether Par3 and Pals1 are functionally coupled, and specifically, whether Par3 is required for the formation of the VMZ. To this end, Par3 or Pals1 expression was silenced in MDCK-II cells using specific siRNAs. Two siRNAs against each Par3 and Pals1 were designed, all of which efficiently reduced the amounts of the respective proteins as determined by immunoblotting (data not shown). We then used semi-quantitative immunoblotting and confocal microscopy to analyze MDCK-II monolayers with reduced Par3 or Pals1 levels (Figure 2B-2I). Loss of Pals1 caused a concomitant loss of PatJ (Figure 2C and 2D), as expected [5], and led to a notable but incomplete reduction in Par3 levels (Figure 2C and 2E). Loss of Pals1 also reduced the amount of TJ-associated Lin7c, while its localization to the lateral membrane appeared unaffected (Figure 2C and 2F). Interestingly, loss of Par3 profoundly reduced TJ localization and total protein levels of Pals1, PatJ and Lin7c (Figure 2C and 2G-2I). Par6 levels were slightly reduced in both Par3- and Pals1-depleted cells, whereas the expression of aPKC, ZO-2, and occludin was largely unaffected (Figure 2C). We concluded that Par3 is critical for the stable expression and localization of Pals1, PatJ and Lin7c, and hence controls, directly or indirectly, the formation of the VMZ.

**Figure 2:**
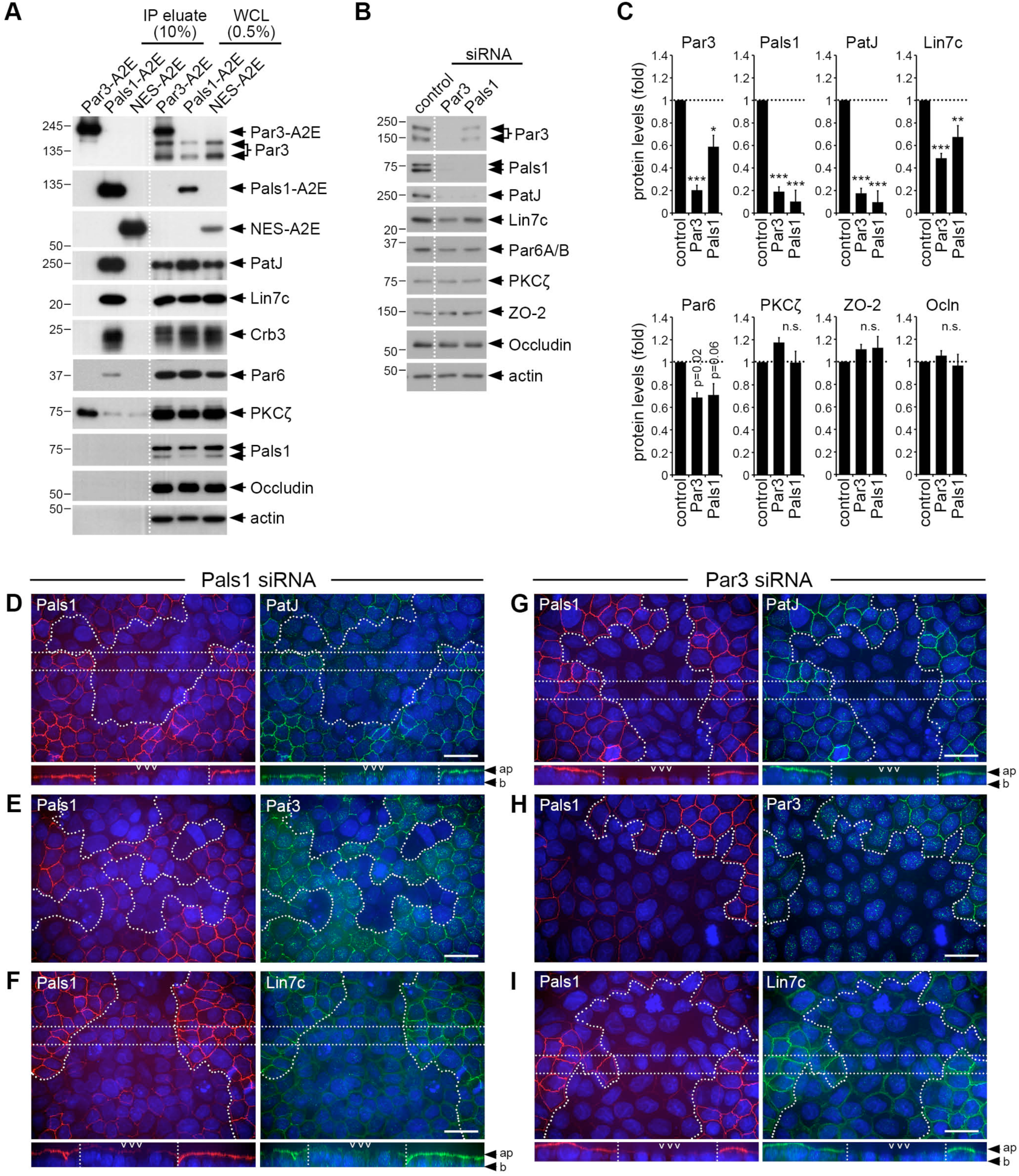
Pals1 forms a distinct protein complex whose stability is dependent upon Par3 expression. **A.** Immunoprecipitations of Par3-A2E, Pals1-A2E and A2E-NES from filter-grown MDCK-II cell lines using anti-GFP antibodies. Membranes were probed with the indicated antibodies. A representative of three independent experiments is shown. **B.** siRNA-mediated knockdown of Par3 or Pals1 in MDCK-II cells. Cells were transfected with 100 nM siRNA against canine Par3 or Pals1, lysed and analysed two-three days post-transfection. Lysates were blotted and probed for the indicated antibodies. **C.** Quantification of data shown in B using densitometry. Error bars show s.e.m. *** = p<0.0001, ** = p<0.001, * = p<0.005, n.s. = not significant, student’s t-test (n=5). **D-I.** Confocal micrographs of siRNA-transfected MDCK-II cells grown on filter membranes. Cells were transfected with Par3 or Pals1 siRNAs, trypsinized two days post-transfection and seeded onto filter membranes. Cells were fixed two days post-seeding and stained with the indicated antibodies and DAPI. Maximum intensity projections of X/Y confocal stacks are shown. X/Z projections were generated from sub-volumes indicated by the vertical lines in the X/Y projections. Ap: apical; b: basal. Scale bars are 20 µm.

### QPP of a Par3-Pals1 pair identifies junction-associated proteins

We designed experiments to test whether pairwise QPP of Par3-A2E and Pals1-A2E would resolve the molecular and spatial organization of the apical-lateral border. To our surprise, published proximity labeling protocols [39] failed to induce biotinylation in fully polarized Pals1-A2E and Par3-A2E MDCK-II cells grown on filter membranes (Figure S4A). We therefore established a modified labeling protocol, which permitted robust and spatially restricted proximity biotinylation in this cell culture system (Figure 3A and 3B, and Figure S4B and S4C). For SILAC-based QPP, Par3-A2E and Pals1-A2E cells were adapted to heavy and light SILAC medium (Par3[H] and Pals1[L], respectively) and compared in a pairwise fashion (Figure 3C). Five independent biological replicates were analyzed, resulting in the identification of ∼1500-1900 proteins per replicate (Figure S4D and S4E). To produce a high-confidence proteome (HCP) of the apical-lateral border, we only considered proteins identified in all five replicates, resulting in a proteome of 1186 proteins (Figure 3D and Suppl. File S1). Importantly, the measured SILAC ratios were remarkably well correlated across all replicates (Figure 3E and 3F), and thus should provide a quantitative and reliable readout to dissect the spatial organization of the HCP along the apico-basal axis (see below).

**Figure 3:**
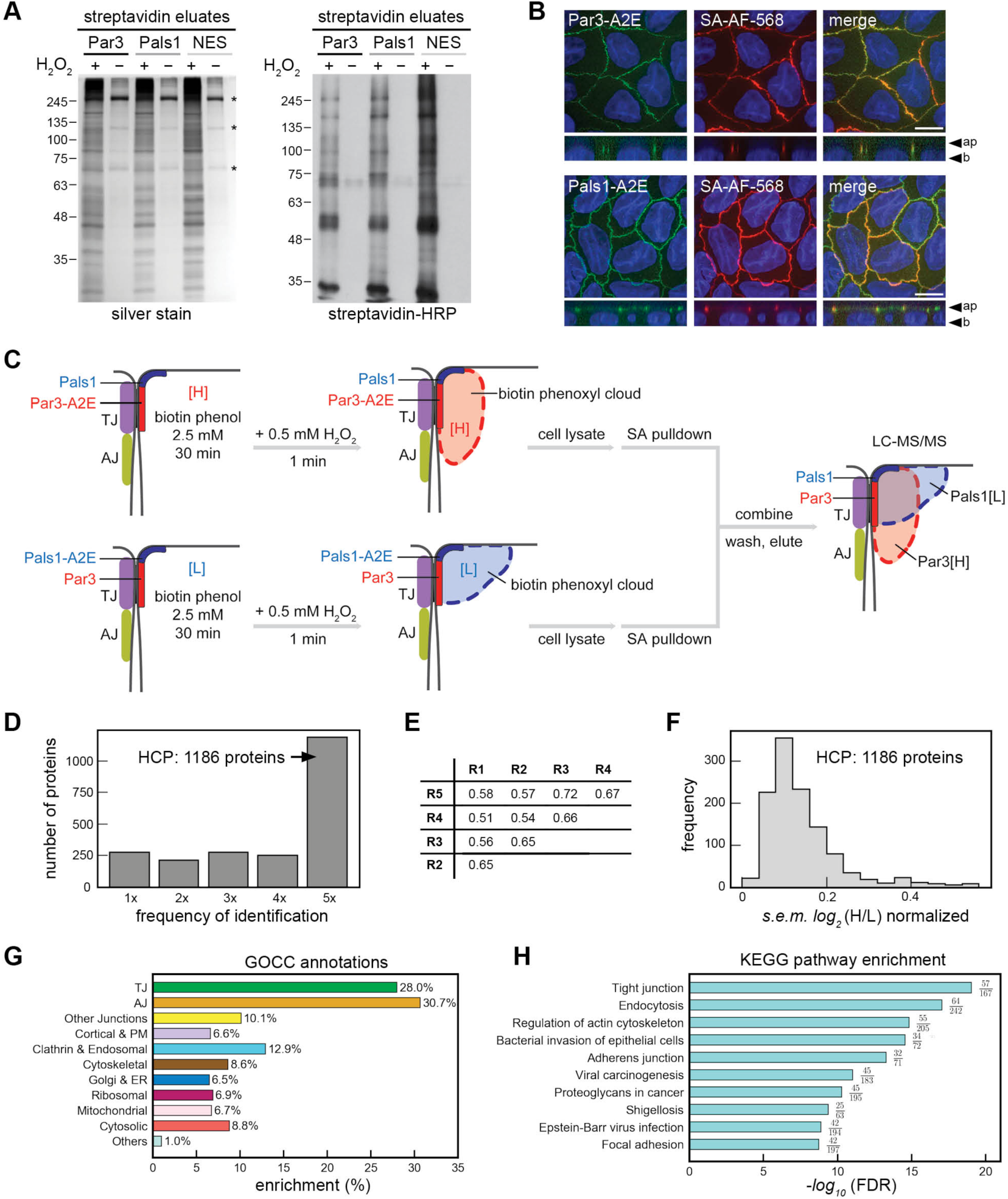
Proximity proteomics of a Par3-Pals1 APEX2 SILAC pair. **A.** Proximity labeling in the Par3-A2E, Pals1-A2E, and A2E-NES cell lines. Cells were pre-incubated with 2.5 mM biotin phenol for 30 min and labeled by the addition of 0.5 mM H_2_O_2_ for 1 min. Biotinylated proteins were purified using streptavidin sepharose, eluted, separated by SDS-PAGE, and either silver-stained (left) or blotted and probed with streptavidin-HRP (right). **B.** Confocal micrographs of filter-grown Par3-A2E and Pals1-A2E cell lines after proximity biotinylation as in A. Cells were fixed and stained with fluorescently labeled streptavidin (SA-AF-566). Maximum intensity projections are shown. Scale bar is 10 µm. **C.** Schematic of pairwise proximity proteomics of Par3-A2E and Pals1-A2E. The anticipated extent of biotinylation generated by Par3-A2E and Pals1-A2E at apical junctions is depicted. **D.** Histogram showing the frequency of protein identifications in the five replicates. Proteins identified in all five replicates represent a high-confidence proteome (HCP) of the apical-lateral border. **E.** R^2^ values of the *log_2_*(H/L) ratios between all five replicates. **F.** Frequency distribution of standard error means (s.e.m.) of *log_2_*(H/L) ratios in the HCP. **G.** Composition of the HCP according to GOCC annotations and a hierarchical categorization regime (see Materials and Methods). **H.** KEGG pathway analysis of the HCP. Pathways are ranked according to false discovery rate (FDR). See also Figure S4.

To evaluate the molecular and functional composition of the HCP we used a combination of Gene Ontology Cellular Component (GOCC) annotations, manual annotations, and KEGG pathway enrichment analyses (see Materials and Methods). As expected, the HCP was highly enriched in TJ and AJ proteins, with approximately 30% of all annotated TJ and AJ proteins being present in the HCP (Figure 3G and 3H). Considering that MDCK-II cells express only 90 out of a total of 124 known TJ proteins, and only 95 out of a total of 130 known AJ proteins [40], the actual enrichment was even higher (∼40% for both TJ and AJ proteins). Moreover, the HCP covered 54% (206/383 proteins) of a previously reported ZO-1 proteome [23], but only 33% (87/263) of a previously reported E-Cadherin proteome [25] (Figure S4F). Thus, the HCP is biased towards TJ components, as expected. Additional protein categories identified in the HCP included other junctional proteins, cortical, plasma membrane (PM), endocytic and cytoskeletal proteins, as well as ER, Golgi, ribosomal, mitochondrial, and cytosolic proteins (Figure 3G). Manual evaluation of the HCP revealed that a large number of proteins annotated as ‘cytosolic’ are signaling proteins (kinases, phosphatases, proteases, proteins regulating ubiquitination, proteins involved in Rho/Ras signaling, adaptors and scaffolds, and transcriptional regulators) and metabolic enzymes (Figure S4G and Suppl. File S2). We concluded that QPP with a Par3-Pals1 APEX2 pair reproducibly identifies a junctional proteome that encompasses a large number of novel junction-associated cytoskeletal, trafficking and signaling proteins.

### QPP of a Par3-Pals1 pair resolves the molecular and spatial organization of the apical-lateral border

To evaluate whether the identified proteome components are spatially resolved, the HCP was displayed as a scatter plot, in which the average peptide intensities were plotted over the mean *log_2_*(H/L) ratios for each protein identified. We then determined the map localization of known apical and lateral membrane proteins and TJ and AJ components using GOCC annotations and information from the primary literature (Figure 4A). We expected that proteins associated with Par3 should be enriched with *log_2_*(H/L) >1, whereas those associated with Pals1 should be enriched with *log_2_*(H/L) <-1. In addition, because the VMZ and TJ are juxtaposed (and possibly continuous with one another), TJ proteins should be shared between Par3 and Pals1, and thus exhibit relatively neutral *log_2_*(H/L) ratios. Indeed, we observed Par3 and Pals1 with *log_2_*(H/L) of at least 2 and −2, respectively (Figure 4A). In addition, PatJ and Lin7c clustered with Pals1, as expected, and so did proteins associated with the apical membrane (e.g. MUC1, EZR, RDX, MSN). By contrast, most bona-fide AJ proteins (catenins, CDH1, CDH6, NECTIN2, NECTIN3) and basolateral membrane proteins (e.g. MET, ITGA2, ITGA3, ITGB1) exhibited *log_2_*(H/L) ratios >0.5. Bona-fide TJ proteins showed *log_2_*(H/L) ratios of −1 to 0.5, and thus were clearly separated from apical, AJ, and lateral proteins. Interestingly, closer examination of our proximity map revealed an apico-basal gradient of TJ proteins (Figure 4B). OCLN (occludin), CGN (cingulin), ZO-1, and ZO-3 occupied an apical position, while CXADR, F11R (JAM-A), MAGI1, MAGI3, CLDN2, and CLDN4 occupied a basal position. ZO-2 and CLDN3 were found in between apical and basal TJ proteins. As expected, the PAR module (PARD3B, PRKCI (aPKC), YWHAZ (Par5), MARK2 (Par1b)) and the SCRIBBLE module (SCRIB, LLGL2, LLGL1) clustered with TJ proteins, with LLGL1 showing a more apical localization than SCRIB and LGLL2. We further noted that PARD6B localized apically of TJ, and that AFDN (afadin) and VCL (vinculin) clustered with basal TJ proteins. In conclusion, our proximity proteome resolves components of TJ and AJ and corroborates that Pals1, PatJ and Lin7c form a protein complex apically of TJ. In addition, the data suggest that TJ are composed of apical and basal subdomains.

**Figure 4:**
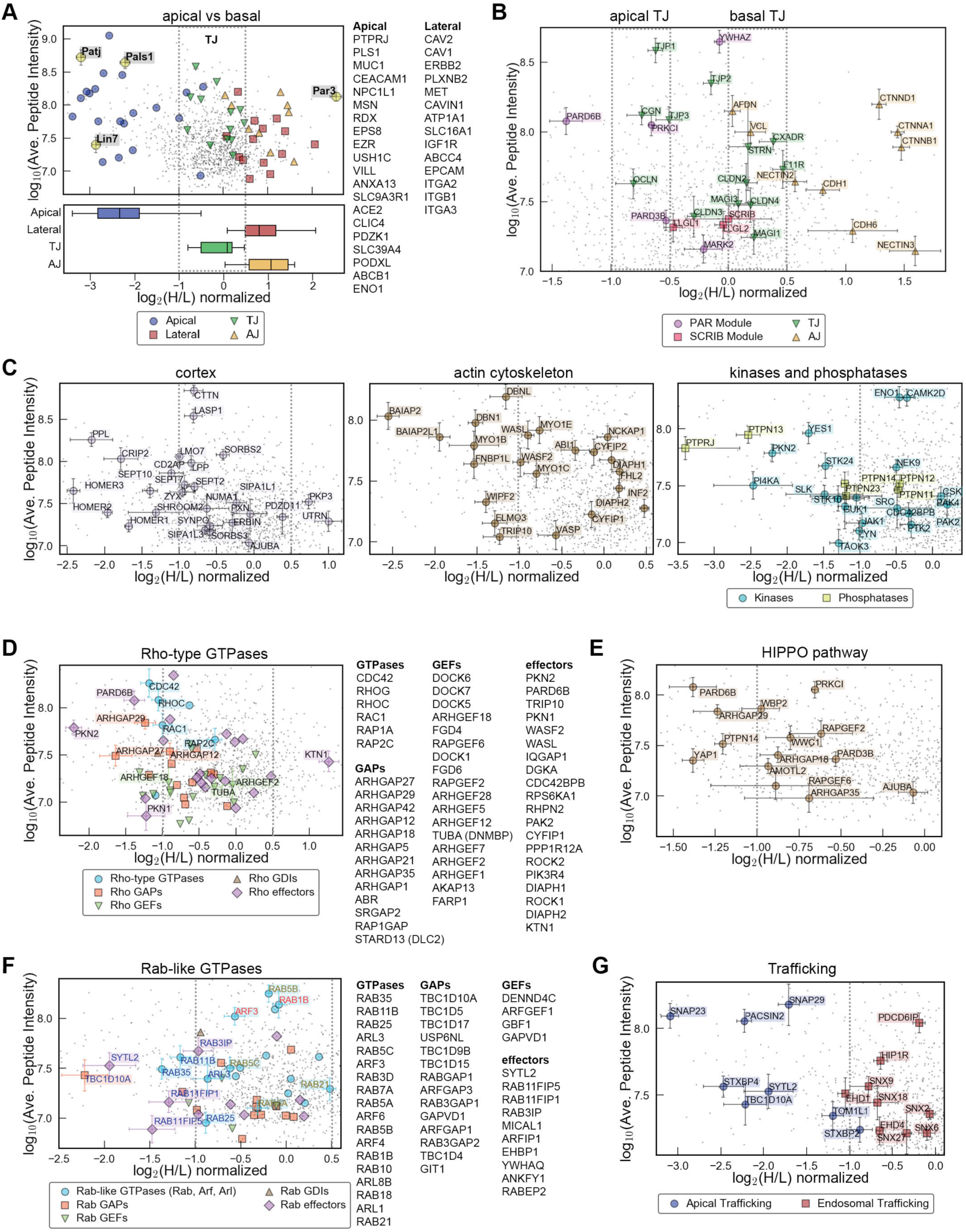
Spatial and molecular organization of the apical-lateral border. **A-G.** Scatter plots showing the average *log_10_* peptide intensities plotted over the average *log_2_*(H/L) normalized SILAC ratios for all proteins identified. Error bars indicate s.e.m. (n=5). **A.** Distribution of apical and baso-lateral proteins as listed on the right. The box plot shows the median distribution of the indicated categories. Apical, TJ and AJ/lateral compartment markers exhibit significantly different distribution profiles (p<0.0001 for all combinations). **B.** Zoom showing the sub-junctional distribution of TJ, AJ, and polarity proteins. Note that TJ proteins form distinct apical and basal protein modules. **C.** Distribution of select cortical, actin and signaling proteins. **D.** Distribution of all Rho-type GTPases, their GEFs and GAPs and effector proteins. **E.** Distribution of select HIPPO pathway proteins. **F.** Distribution of all Rab-like GTPases, their GEFs and GAPs and effector proteins. **G.** Distribution of select trafficking regulators. Proteins in A, D and F are listed in ascending *log_2_*(H/L) ratios (i.e. apically localized proteins are on top). See also Figure S5.

To facilitate access to our proteomics data we have generated an interactive online portal that permits the localization of proteins and protein categories identified in our Par3-Pals1 proteome to be displayed in the scatter plot (Suppl. File S3). Figures 4C-4G show the map localization of select cortical and actin-associated proteins, signaling molecules, small GTPases, and exocytic and endocytic trafficking regulators. This data can be summarized as follows: Firstly, certain cortical proteins (e.g. HOMER proteins), regulators of the actin cytoskeleton (e.g. BAIAP2, BAIAP2L1, FNBP1L), and signaling molecules (e.g. PTPN13, PKN2, STK24, YES1) with unknown or poorly defined subcellular localizations exhibited pronounced negative *log_2_*(H/L) ratios (Figure 4C). Such proteins are likely to be associated with the VMZ or with the apical cortex. Secondly, we find that the repertoire of TJ-associated small Rho-type GTPases and their corresponding GEFs and GAPs is much larger than previously recognized (Figure 4D) [41]. Known GEFs and GAPs of the AJC included ARHGEF18 (p114 RhoGEF) and ARHGAP12, which clustered with the apical TJ pool, and ARHGEF2 (GEF-H1) and TUBA (DNMBP), which occupied a more basal TJ location. Potentially novel regulators of the apical-lateral border included several DOCK proteins, ARHGAP27, ARHGAP42, ARHGEF1, ARHGEF5, and ARHGEF28. In addition, we identified ARHGAP29, ARHGAP18, ARHGAP5, ARHGAP35, RAPGEF2, and RAPGEF6, all of which have been implicated in HIPPO/YAP signaling [42–45] (Figure 4E). In total, the HCP contained 58 (out of an estimated 110) conserved HIPPO pathway proteins [46]. This underscores the role of TJ, and in particular the apical polarity proteins, in the regulation of small GTPase and HIPPO signaling. Thirdly, proteins involved in apical trafficking (e.g. RAB11B, RAB11FIP1, RAB11FIP5) and apical membrane fusion (e.g. SNAP23, SYTL2) exhibited markedly more negative *log_2_*(H/L) ratios than proteins involved in early endosomal transport (e.g. RAB5A, RAB5B, sorting nexins) (Figure 4F and 4G). Potentially novel regulators of the apical trafficking machinery included TBC1D10A, STXBP4, and PACSIN2. Taken together, this analysis showed that the Par3-Pals1 proximity map resolves the polarized organization of the apical-lateral cortex as well as the epithelial signaling and trafficking machinery.

Lastly, we addressed the significance of ER, ribosomal, and RNA-binding proteins identified in the HCP (Figure S5A). RNA-binding and ribosomal proteins showed relatively neutral *log_2_*(H/L) ratios and are common contaminants in AP-MS experiments [47]. Hence, whether such proteins are specifically associated with the AJC, as supported by a previous study [48], remains to be determined. By contrast, ER residents exhibited pronounced positive *log_2_*(H/L) ratios, suggesting that the ER is located in close proximity to the basal aspect of the AJC. Indeed, EM supported this notion and further indicated an ‘ER-free zone’ near the apical cortex and TJ (Figure S5B). Whether the ER (and perhaps ER-PM contacts) are specifically enriched at the basal aspect of the AJC while being excluded from other areas of the lateral cell cortex requires a more detailed analysis.

### A spatially resolved TJ-associated protein interaction network

To analyze the physical connectivity of the HCP we retrieved PPIs from the BioGRID database [49, 50] und used Cytoscape [51] to illustrate and analyze the topology of the resulting network. Entries based on previous TJ and AJ proteomes were excluded to avoid network connections that may not be based on direct PPIs. This produced a highly interconnected network of 1127 proteins (nodes) and 11934 interactions (edges) (Figure S6A-S6C). The remaining 59 proteins could not be mapped due to the lack of annotated binding partners within the network. This included known components of the apical-lateral border, such as EPCAM, SYTL2, CXADR, ARHGEF18 and ARHGAP18. In order to simplify visualization and interpretation of the interactome we eliminated chaperones, ER- and ribosomal, RNA-binding, and metabolic proteins, resulting in a core AJC-associated network of 807 proteins and 6498 interactions (Figure S6D and S6E). The topology of this network, analyzed using the network analyzer [52] in Cytoscape, followed the power law typical of biological systems [53] (Figure S6F), with a clustering coefficient of 0.21, an average number of neighbors of 16.1, and a characteristic path length of 2.78 (Figure S6G). These parameters indicate that the AJC-associated network is denser than the previously described E-Cadherin interactome [26] and similarly dense than the protein network of mitochondria [54].

To unravel the spatial organization of the network, nodes were spread out in two dimensions (Figure 5). The measured *log_2_*(H/L) ratios were used to resolve the apico-basal organization of the proteome, while subcellular localization and molecular function inferred from manual annotation of the HCP (Figure S4G) was applied as a ‘spatial ruler’ for membrane-proximity. We further defined membrane domain boundaries using TJ proteins and the polarity proteins as spatial landmarks (Figure 4B). OCLN and F11R demarcated the apical and basal boundaries of the TJ (*log_2_*(H/L) −1 to 0.5), thereby separating components of the apical compartment (Pals1, PatJ, Lin7c, PARD6B) and the lateral membrane (catenins, cadherins, nectins). This network representation demonstrated that membrane-proximal and membrane-distal, as well as apical and basal components of the HCP were highly interconnected. Apical and basal TJ proteins formed a core TJ network of 606 proteins and 4202 interactions (Figure S6H). This network exhibited topological characteristics similar to the entire network (Figure S6G). We concluded that the HCP is composed of a core of TJ-associated proteins that are physically connected to components of the apical cortex and AJ.

**Figure 5:**
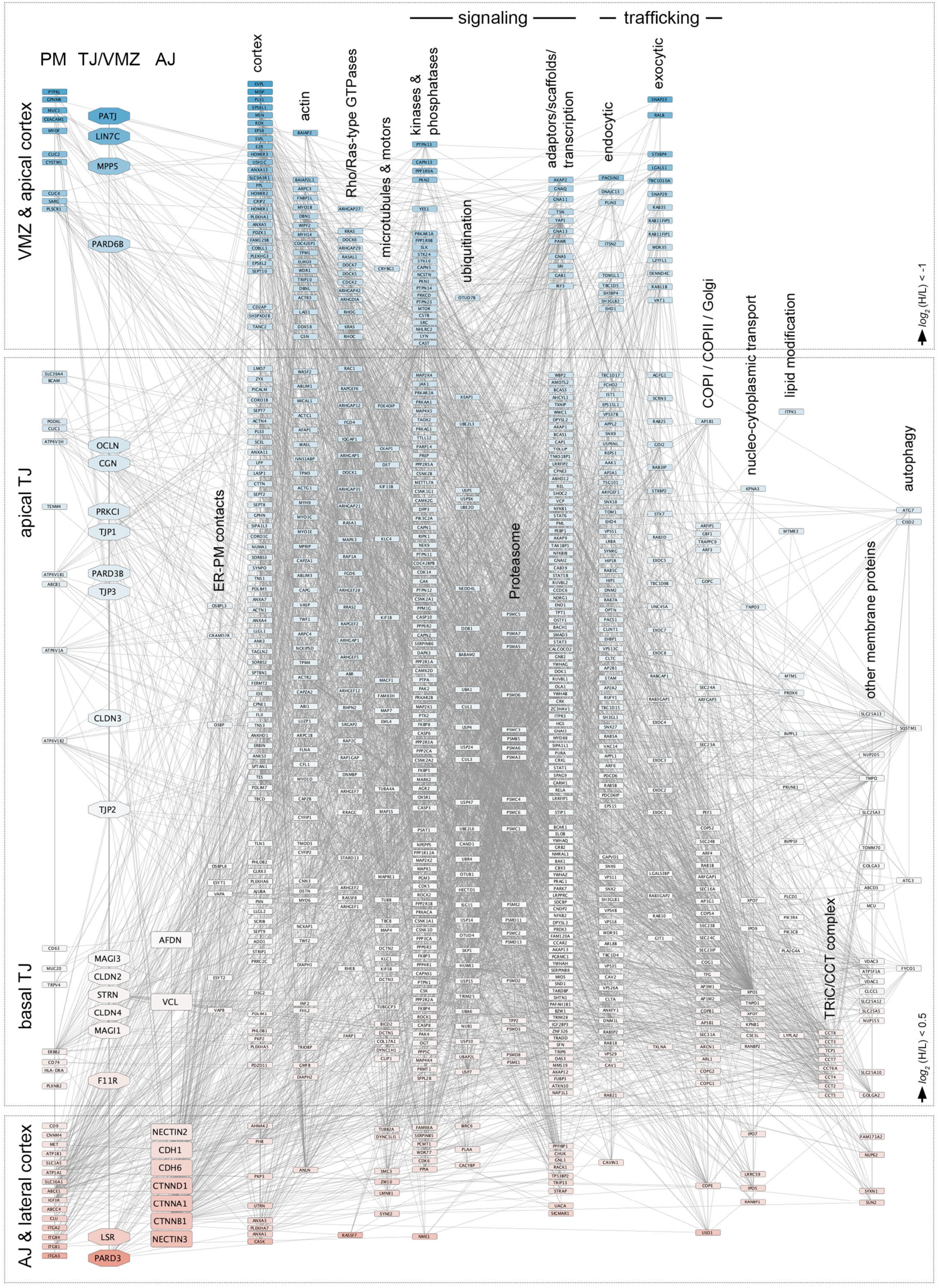
A spatially resolved protein interaction network of the apical-lateral border. Shown is a PPI network of the HCP (807 proteins). Interactions were retrieved from the BioGRID database. ER, ribosomal, and RNA binding proteins, chaperones, metabolic, and unknown proteins were excluded from the interactome. Proteins were ranked top to bottom according to their *log_2_*(H/L) ratios, and from left to right according to their anticipated proximity to the membrane. Domain boundaries were defined based on the *log_2_*(H/L) ratios of integral AJC components and polarity proteins. Gaps in between domain boundaries are exaggerated for the sake of clarity. See also Figure S6.

### Defining the VMZ-associated protein network

Our data so far strongly suggested that Pals1 and its known interactors define a VMZ apical of TJ. To identify proteins directly associated with the VMZ, a pairwise SILAC experiment was conducted in which the Pals1-A2E cell line was compared with a cell line expressing APEX2 in the cytoplasm (A2E-NES) (Pals1-A2E [H] vs A2E-NES [L]). We reasoned that cytoplasmic APEX2 should greatly reduce ‘cortical and membrane-distal bystanders’ and hence produce a more specific Pals1 proximity proteome (Pals1 PP) enriched in proteins that are physically connected to the VMZ. An identical experiment was performed with the Par3-A2E cell line (Par3-A2E [H] vs A2E-NES [L]) (Figure 6A). 117 and 99 proteins were significantly enriched with Pals1-A2E and Par3-A2E, respectively (see Materials and Methods) (Suppl. Files S4 and S5). As expected, the Pals1 and Par3 PPs were markedly different in composition; only 23 proteins were common to both proteomes (Figure 6B). Pals1, PatJ, Lin7c as well as HOMER3, SNAP23, and ARHGAP29 were all highly enriched in the Pals1 PP, whereas Par3, RASSF7, and certain signaling and trafficking proteins (e.g. VPS13A, SORL1, PUM2) were specifically enriched in the Par3 PP (Figure 6C and 6D). Moreover, the Pals1 and Par3 PPs occupied distinct locations in the Par3-Pals1 scatter plot (Figure 6E). While the Pals1 PP was positioned precisely in between components of the apical cortex and TJ, the Par3 PP encompassed proteins of a more basal compartment, including TJ and AJ proteins (Figure 6D). Of note, ER and ribosomal proteins were also abundant in the Par3 PP, again supporting the view that the basal aspect of the AJC, and perhaps Par3 directly, is associated with the ER and the protein translation machinery. Interactomes for the Pals1 and Par3 PPs generated based on BioGRID entries confirmed that Par3 and Pals1 are linked to different protein networks (Figure 6F and 6G). We focused on the Pals1 PP and found that the VMZ core complex (PatJ, Pals1, Lin7c, Par6) was physically associated with the scaffolding proteins HOMER3 and HOMER1, the membrane fusion regulator STXBP4, and the HIPPO pathway protein NF2 (Merlin) (Figure 6F). In addition, the phosphatases PTPN13 and PTPN21, the serine-threonine kinase and Rho/Rac effector PKN2 (PRK2), and the RhoGAP ARHGAP29 were associated with the VMZ via STXBP4. SNAP23, MUC20, and a number of other proteins found in close proximity to Pals1 could not be mapped into the network. TJ proteins formed a separate sub-network, which may be linked to the VMZ via interactions between PatJ and ZO-3 [6]. Apical membrane proteins, a hub of apical cortical proteins, actin regulators and trafficking proteins, and proteins involved in small G protein signaling constituted additional clusters.

**Figure 6:**
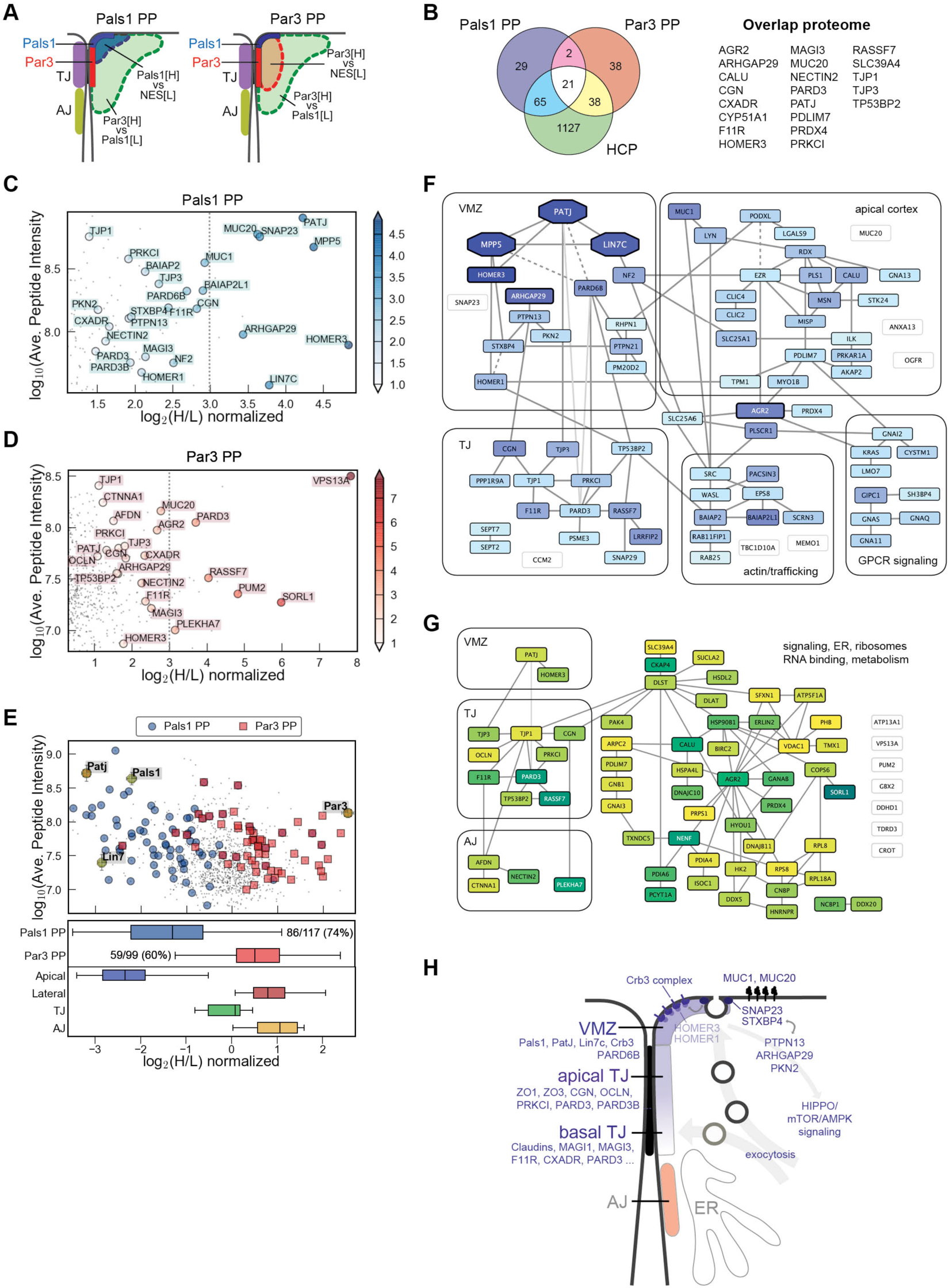
An interactome of the VMZ. **A.** Schematic illustrating the anticipated extent of proximity labeling produced by a Pals1-NES and Par3-NES SILAC pairs compared to the Par3-Pals1 SILAC pair. **B.** Venn diagram showing the degree of common and unique components of the Par3 and Pals1 PPs compared to the HCP. Note the limited overlap between the Par3 and Pals1 PPs. Proteins identified in all replicates are shown. **C.** Scatter plot highlighting select proteins identified in the Pals1 PP. **D.** Scatter plot highlighting select proteins identified in the Par3 PP. **E.** Scatter plot and box plot showing the distribution of the Par3 and Pals1 PPs in the Par3-Pals1 proteome. Apical, TJ, AJ, and lateral proteins are shown as spatial reference. **F.** Interactome of the Pals1 PP drawn in Cytoscape. PPIs could be retrieved for 78 out of the 117 proteins present in the Pals1 PP. Protein nodes are colour-coded according to their *log_2_*(H/L) ratios in the Pals1 PP. Dark blue nodes with white font exhibited *log_2_*(H/L) > 3. Proteins shown in white nodes were identified with *log_2_*(H/L) > 2 but were not connected to the network. PPIs shown as dashed lines were manually added: PATJ-PARD6B: PMID 19255144, MPP5-PARD6B: PMID 15140881, HOMER1-STXBP4: BioGRID pre-pub entry, PODXL-EZR: PMID 11461930. Interactions between PARD3 and PATJ and PARD3 and PARD6B are shown as a light grey line and may not occur in fully polarized MDCK-II cells. **G.** Interactome of the Par3 PP drawn in Cytoscape. PPIs could be retrieved for 67 out of the 99 proteins present in the Par3 PP. Protein nodes are colour-coded according to their *log_2_*(H/L) ratios in the Par3 PP. Dark green nodes with white font exhibited *log_2_*(H/L) > 3. Proteins shown in white nodes were identified with *log_2_*(H/L) > 3 but were not connected to the network. **H.** Model of the organization of the apical-lateral border and possible functions for the VMZ. See also Figure S7.

We sought to confirm that the VMZ-associated proteins identified here indeed form a PPI network at the apical-lateral border. PTPN13 and PKN2 were previously shown to localise to TJ and to be involved in TJ assembly and epithelial barrier regulation [55, 56]. It is also known that PTPN13 interacts with PKN2 and ARHGAP29 via PDZ domain interactions [57, 58] (Figure S7A). To test whether ARHGAP29, HOMER proteins, and STXBP4 localize to apical junctions, we transfected the respective cDNAs into MDCK-II cells. Indeed, HA-ARHGAP29, HA-HOMER3, and HA-HOMER2 efficiently colocalised with Pals1 by confocal microscopy. HA-STXBP4 produced a diffuse cytoplasmic staining and only partially colocalized with Pals1 at the membrane (Figure S7B). In conclusion, we show that the VMZ is defined by a stable protein complex composed of Pals1, PatJ, Lin7c and Crb3, which connects Par6, HOMER proteins, ARHGAP29, STXBP4, and a specific set of kinases and phosphatases to the apical-lateral border.

## Discussion

Compartmentalization of the cell cortex is critical for epithelial form and function and generates signaling platforms and structural scaffolds for the spatio-temporal control of the epithelial polarity and morphogenesis program. However, the precise nanometer-scale organization of the epithelial cell cortex is not well defined. Here we have produced the first quantitative spatio-molecular map of the apical-lateral border in mammalian epithelial cells, and in doing so, uncover an unexpected degree of sub-junctional compartmentalization. In essence, we show that the mammalian AJC is composed of at least four membrane compartments – the VMZ, apical and basal TJ, and AJ (Figure 6H). We identify hundreds of potentially novel structural and regulatory components of the AJC and define in detail the composition of the VMZ-associated proteome. We provide evidence that our proteome is instructive in predicting a protein’s sub-junctional localization, and possibly, in identifying functional relationships between spatially correlated proteins. Hence, APEX2-mediated EM and QPP quantitatively dissects the architecture of cell membranes at the nanometer level. Combined with multiplex proteomics and a larger repertoire of APEX2 fusion proteins this opens the way for a comprehensive spatio-molecular analysis of the epithelial cell cortex and other subcellular organelles and structures.

Our spatially resolved proteome of the apical-lateral border shares similarities but is not identical to previously published ZO-1 and E-Cadherin proteomes [23, 25, 26]. This is not surprising given that all previous proteomes were generated from cells grown on plastic, a culture system that does not support proper polarity development. Additional factors that are likely to have contributed to the observed differences in proteome components are the different labeling chemistries of BirA* and APEX2, the different bait proteins used, and the different quantification approaches applied. Importantly, previous TJ and AJ proteomes could not unequivocally assign the identified proteins to a specific membrane compartment. By contrast, our approach of using an APEX2 pair of two junction-associated proteins allowed us to quantify the relative sub-junctional localization of known and novel components of the apical-lateral border. Not only did this approach resolve TJ and AJ and demonstrate that Pals1 defines a distinct membrane domain apical of TJ; our data also provide evidence for further sub-compartmentalization of TJ along the apico-basal axis (Figure 6H). TJ proteins such as occludin, cingulin, ZO-1, and ZO-3 formed an apical TJ domain, which was clearly separated from a basal TJ domain composed of claudin-2, claudin-4, MAGI-1, MAGI-3, F11R, and CXADR. Interestingly, this bimodal organization is consistent with a recent report showing that claudin-2 and claudin-4 are incorporated into TJ from the basal end, while occludin enters TJ from a more apical location [59]. They also provide a molecular basis for the ultrastructural organization of TJ strands observed by freeze-fracture EM, in which basal strands appear more loosely organized than apical ones, which has long been suspected to be due to compositional and perhaps functional differences between apical and basal TJ [60–62]. The combined data support and extend upon a model recently proposed by Van Itallie et al [59], in which the basal aspect of TJ receives incoming integral membrane proteins of TJ (such as claudins, F11R and CXADR) and incorporates these into nascent TJ strands. Delivery may be mediated by vesicular transport and controlled by the exocyst complex [63–65]. Basal strands then mature into apical strands, in which claudins and occludin become linked to the cytoskeleton via ZO proteins. Interestingly, claudin-3 showed a more apical TJ localization in our proximity map than claudin-2 and claudin-4. This is in line with data showing that different claudins are targeted to and incorporated into TJ with different kinetics [59], and that at least in certain cell types, claudins can be concentrated at different apical and basal domains at steady state [66]. Taken together, our data indicate that TJ are composed of two subdomains. Such a modular organization may have important implications for TJ formation, maturation and plasticity, and for the control of the various cellular processes that are mediated by this multifunctional membrane compartment.

We further show that Pals1, PatJ, Lin7c and Crb3 form a stable protein complex, whose formation is strictly dependent upon Par3 expression. EM imaging and QPP indicate that this protein complex defines the margins of the apical membrane, that is, a membrane domain apical of Par3, and hence apical of TJ. These observations prompt us to propose that the VMZ identified here represents the mammalian counterpart of the invertebrate MZ. This implies that the molecular and spatial organization of the apical polarity proteins is much more conserved than previously recognized, and argues against previous speculations that the MZ is an evolutionary precursor of TJ [10, 11]. A notable difference between the invertebrate MZ and the VMZ is that the former is located at the apical-most position of the lateral membrane, whereas the latter is located at the margins of the free apical membrane. A possible explanation for this differential localization may be found in the evolution of the Crb protein family. Invertebrates possess only one Crb protein, which localizes to the MZ and has been shown to mediate cell-cell adhesion via homophilic interactions between its large extracellular domains [67]. By contrast, vertebrates encode three Crb orthologs. With the exception of epithelia in the brain, cornea and retina (which express Crb1 and Crb2), all other epithelia express Crb3. Crb3 lacks the bulk of the extracellular domains of Crb1 and Crb2 and hence is unlikely to be involved in cell adhesion. It is tempting to speculate, therefore, that with gene duplications of Crb during vertebrate evolution, the extracellular domain of Crb3 became dispensable for Crb3 function, causing the MZ to lose its cell-cell adhesion function and to ‘move’ apically. This hypothetical event is likely to have occurred concomitantly with (or may even have been the consequence of) the reorganization and regrouping of integral junctional components and the advent of TJ [10, 11].

What then, could be the function of the VMZ? It is well established that the Crb module acts upstream in the HIPPO pathway, both in invertebrates [68–70] and vertebrates [71–73], and that HIPPO signaling output is regulated by mechanical and cytoskeletal cues [74]. In addition, Crb3, Pals1 and PatJ are all required for TJ formation and epithelial polarity development [6, 7, 18]. Interestingly, the Crb module has also been implicated in mTOR/AMPK signaling [75, 76]. HIPPO and mTOR/AMPK signaling pathways are interconnected [77, 78] and tightly coupled to TJ formation and polarity development [79]. The molecular machinery that integrates these processes downstream of the Crb module is not well defined. We identify here a VMZ-associated interactome of proteins with reported or proposed functions in HIPPO signaling (ARHGAP29, STXBP4, PTPN13) [44, 45, 80, 81], TJ formation (PKN2 and ARHGAP29) [55, 82, 83], cell polarity (HOMER3) [84], and mTOR/AMPK signaling (PKN2) [85, 86]. We speculate that this protein network receives mechanical and nutrient sensing ques at the VMZ and transmits these to the HIPPO and mTOR/AMPK signaling pathways. Furthermore, we find that SNAP23 is associated with the VMZ. SNAP23 can form an apical SNARE complex with STX3 [87, 88], but can also assemble into a ternary complex with STX4 and STXBP4, in which STXBP4 acts as a negative regulator of membrane fusion [89]. This might suggest an additional role for the VMZ in regulated exocytosis. Indeed, a function for Pals1 in polarized membrane trafficking has been reported previously [90, 91], and it is well established that membrane proteins are delivered to the apical surface at a site juxtaposed to TJ [92, 93], a location that coincides with the position of the VMZ identified here. Taken together, we propose that the VMZ serves as a specific membrane scaffold that links nutrient sensing and mechanical signals to the control of membrane trafficking, polarity, growth, and survival [74, 94, 95] (Figure 6H). How the here described VMZ-associated interactome integrates these processes remains to be determined.

## Acknowledgements

We thank Andre le Bivic, Spiros Efthimiopoulos, Ben Margolis, Karl Matter, Jingshi Shen, and Ceniz Zihni for providing antibodies, plasmids and other reagents or tools. This work was supported by research grants to SS (RG39/14 and NIM/03/2016), an NTU start-up-grant to AL, and funding by the Agency for Science and Technology (A*STAR), Singapore, to WH and JG. BT was the recipient of a PhD scholarship from the A*STAR Graduate Academy. We further thank the NTU Institute of Structural Biology (NISB) for support.

## Declaration of Interest

The authors declare no conflict of interest.

## Author contributions

BT conducted experiments and analyzed the data. SP conducted the MS analysis. AL designed and conducted experiments, analyzed the data, and wrote the manuscript. JG, WH, and SS designed experiments.

## Materials and Methods

### DNA constructs

pcDNA3-mito-APEX was purchased from Addgene (Plasmid #42607) and the A134P point mutation was introduced to convert APEX into APEX2 [32]. APEX2 was subsequently inserted into Clontech N1 and C1 vectors to produce parental vectors (N1-A2E and C1-A2E, respectively). Pals1 cDNA (NM 022474, Image Clone 30520451) was purchased from Source Bioscience and cloned into the N1-A2E vector using XhoI and SacII sites, producing Pals1-A2E. Par3 cDNA (HA-Par3, NM 001184785) was provided by Ben Margolis and subcloned into the N1-A2E vector using BglII and HindIII, producing Par3-A2E. Occludin cDNA (NM 002538) was subcloned into the C1-A2E vector using the SalI and SacII sites. Nuclear export signal (NES) sequence (5’-CGCTGCAGCTGCCTCCCCTGGAGCGCCTGACCCTGGACTAATAG-3’) was purchased as a pair of complementary oligonucleotides with HindIII (5’) and BamHI (3’) overhangs, annealed, and cloned into the C1-A2E vector. HA-ARHGAP29 was purchased from Addgene (Plasmid #104154). cDNAs for STXBP4 and HOMER1, 2, 3 were from Jingshi Shen and Spiros Efthimiopoulos, respectively, and subcloned into C1-3xHA and C1-A2E vectors. All plasmids were verified by sequencing.

### Cell culture and transfection

MDCK-II cells were grown in DMEM (Gibco) supplemented with 10% fetal bovine serum (FBS) and 100 units/ml penicillin, 100 μg/ml streptomycin (Gibco) at 37°C and 5% CO_2_. Cells were seeded onto polycarbonate or polyethylene 0.4 μm-pore Transwell filter inserts at a confluent density of ∼220,000 cells/cm^2^. Media was replenished every two days. Cells were grown on the filter inserts for 7 to 12 days post-seeding to achieve full polarization. To create stable cell lines expressing APEX2-EGFP fusion proteins, MDCK-II cells were transfected with 1 μg plasmid DNA per 10^6^ cells using either polyethlyenimine (Polysciences Inc) or Lipofectamine-2000 (ThermoFisher Scientific). Clonal cell lines were selected with 500 μg/ml geneticin (Gibco) for at least 14 days. All stable lines were subsequently cultured in media containing 200 µg/ml geneticin.

### Immunoprecipitation

MDCK-II cells grown Transwell filters were washed twice in PBS containing 1 mM calcium chloride and 0.5 mM magnesium chloride (PBS^++^). Cells were incubated with cross-linking solution (0.25 mM dithiobis(succinimidylpropionate) (DSP) freshly diluted in PBS^++^ from a 100x DMSO stock) for 15 min at room temperature. The cross-linking was stopped by adding 1 M Tris pH 7.4 stock to a final concentration of 100 mM Tris. Cells were washed once with 150 mM NaCl, 50 mM Tris pH 7.4 and lysed with 0.5% Triton X-100 in 50 mM Tris pH 7.4, 150 mM NaCl, 1 mM EDTA with protease inhibitor cocktail (Roche). Lysates were cleared by centrifugation (20,000x g, 20 min) and incubated with GFP-Trap beads (ChromoTek) for 3 hr at 4°C. Beads were pelleted by centrifugation at 500x g for 2 min at 4°C and washed three times. Proteins were eluted by boiling for 5 min in 4x LDS (Invitrogen) sample buffer containing 200 mm DTT.

### Western blotting

Proteins were separated by SDS-PAGE and blotted onto polyvinylidene difluoride (PVDF) membranes in a wet-transfer set-up at 20 V overnight at 4°C. After blotting, membranes were dehydrated in methanol for 10 s and allowed to dry for at least 1 hr. Membranes were incubated with antibodies diluted in 5% fat-free milk in PBST (PBS, 0.2% Tween-20) or 2.5% bovine serum albumin (BSA), 0.4% Tween-20 in PBS. Membranes were washed three times with PBST, incubated with horseradish peroxidase (HRP)-conjugated secondary antibodies (ThermoFisher Scientific) diluted in 5% fat-free milk in PBST, washed three times with PBST, and developed with chemiluminescent substrate. All antibodies were tested and used at optimal concentrations for WB and IF (see table 2).

**Table 1:**
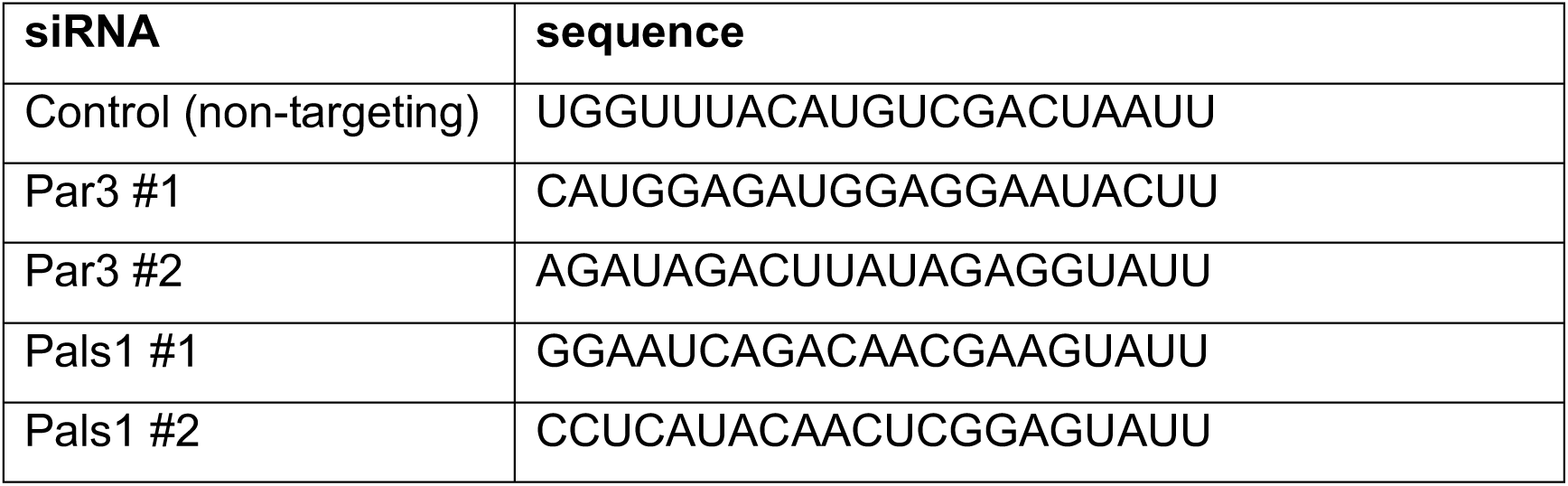
siRNAs

**Table 2:** Antibodies

| Antigen | Company/ Catalog No. | Host Species | Dilution IF | Dilution WB |
| --- | --- | --- | --- | --- |
| Cld1 | Thermofisher/ 71-7800 | Rabbit | 1:500 | 1:5000 |
| Occludin | Invitrogen/ 33-1500 | Mouse | 1:500 | 1:5000 |
| Pals1 | Santa Cruz/ sc-365411 | Mouse | 1:200 | 1:2500 |
| PATJ | LifeSpan Biosciences/ LS-C410011 | Rabbit | 1:200 | 1:5000 |
| Lin7 | Thermofisher/ 51-5600 | Rabbit | 1:500 | 1:10000 |
| Crb3 | Anti-serum/ Ceniz Zihni | Rabbit | - | 1:50 |
| Par3 | Merck Millipore/ 07-330 | Rabbit | 1:500 | 1:5000 |
| PKCζ | Santa Cruz/ sc-17781 | Mouse | - | 1:5000 |
| PKCζ | Santa Cruz/ sc-216 | Goat | - | 1:1000 |
| Par6 | Santa Cruz/ sc-33898 | Goat | - | 1:1000 |
| ZO-1 | Thermofisher/ 33-9100 | Mouse | 1:500 | 1:2500 |
| ZO-2 | Cell Signaling Technology/ 2847 | Rabbit | - | 1:2500 |
| ZO-3 | Invitrogen/ 36-4100 | Rabbit | 1:500 | 1:2500 |
| β-catenin | BD Biosciences/ 610153 | Mouse | 1:1000 | 1:10000 |
| E-cadherin | BD Biosciences/ 610181 | Mouse | 1:1000 | 1:20000 |
| Actin | Sigma Aldrich/ A5060 | Rabbit | - | 1:20000 |
| GFP | Roche/ 11814460001 | Mouse | - | 1:5000 |
| HA | Roche/ clone 3F10, High Affinity | Rat | 1:500 | - |

### Immunofluorescence

MDCK-II cells grown either on glass coverslips or Transwell filter inserts were fixed for IF either by methanol or paraformaldehyde (PFA) fixation. In methanol fixation, cells were washed twice in PBS^++^, once in −20°C methanol, and fixed in methanol at −20°C for 10 min. Cells were washed three times in PBS before blocking. For PFA fixation, cells were washed twice in PBS^++^, fixed in 1% or 3.7% PFA for 15 min or 20 min at room temperature, washed three times in PBS, pre-extracted with 0.5% Triton X-100 in PBS for 10 min, and washed three times in PBS before proceeding. Cells were blocked in 0.2 μm-filtered 10% FBS in PBS for at least 2 hr at room temperature or overnight at 4°C. If cells were grown on Transwell filter inserts, filter inserts were cut out with a sharp razor blade at the end of the blocking step. For staining and washes, filter pieces were laid face-up on parafilm in a humidified chamber and overlaid with 25 μl of staining/wash solutions. Glass coverslips were laid face-down over 50 μl of staining solutions on parafilm in a humidified chamber and were laid face up on parafilm for washes. Samples were incubated with primary antibodies diluted in antibody incubation solution (0.2 μm-filtered 0.1% BSA, 0.01% Tween-20 in PBS) for 1.5 hr at room temperature (see table 2 for antibody dilutions). Samples were washed three times 5 min each in antibody incubation solution and further incubated for 1 hr with fluorescently conjugated secondary antibodies diluted in antibody incubation solution at room temperature shielded from light. Samples were washed three times 10 min each in PBS and stained for 10 min with 20 ng/ml DAPI diluted in PBS. Filter pieces were placed face-up on glass slides, overlaid with a drop of mounting medium (Vectashield, Vector Laboratories H-1000) and covered with glass coverslips. Samples grown on glass coverslips were mounted with a drop of mounting medium. Coverslips were sealed with nail polish and stored at 4°C. Samples were imaged using the CorrSight spinning disk confocal microscope (FEI Company) equipped with an Orca R2 CCD camera (Hamamatsu). Imaging of cells grown on Transwell filters was performed using a 63× oil objective (NA 1.4) (Plan Apochromat M27, Zeiss) and standard filters. Confocal stacks were processed in the FIJI distribution of ImageJ.

### Stable isotope labelling of amino acids in cell culture (SILAC)

SILAC media was prepared by supplementing dialyzed DMEM (ThermoFisher Scientific) with 10% dialyzed FBS (Gibco), 100 U/ml PS, and 0.4 mM isotopically-labelled L-arginine and 0.8 mM isotopically-labelled L-lysine (Cambridge Isotope Laboratory) (“Heavy” media) or the corresponding concentrations of natural isotope compositions of L-arginine and L-lysine (“Light” media). MDCKII cells were metabolically labelled by culturing in SILAC media for at least three passages and verified by MS that they had at least 95% heavy isotope incorporation. For each SILAC experiment, one cell line is grown in “Heavy” media and another in “Light” media on 75 mm Transwell filter inserts.

### Proximity biotinylation and affinity capture

The APEX biotinylation protocol was based on Hung *et. al* [39] with several modifications. MDCK-II cells were grown on either glass coverslips, plastic dishes, or Transwell filter inserts. Cells were incubated with media containing 2.5 mM biotin phenol (Iris Biotech) for 30 min at 37°C. Cells were washed twice with PBS^++^ and the APEX reaction was initiated with 500 μM H_2_O_2_ in PBS^++^. The APEX reaction was quenched by transferring the cells to a quencher solution (PBS^++^ containing 5 mM (±)-6-hydroxy-2,5,7,8-tetramethylchromane-2-carboxylic acid (Trolox), 10 mM sodium ascorbate, 10 mM sodium azide), and washed twice more in the quencher solution. Cells were then either fixed for fluorescence microscopy or lysed in ice-cold RIPA buffer (50 mM Tris pH 8.0, 150 mM NaCl, 5 mM EDTA, 0.1% sodium dodecyl sulphate (SDS), 0.5% sodium deoxycholate, 1% Triton X-100) with protease inhibitor cocktail. Cells were scraped off the plastic dish or the filter insert using a cell scraper. Lysates were briefly sonicated and clarified by centrifugation at 21,500 g for 30 min at 4°C. Protein concentrations were measured using the Bradford’s assay and equalized. For SILAC-MS, biotinylated proteins were incubated with pre-equilibrated streptavidin sepharose beads (GE Healthcare) (50 μl bead slurry per 4 mg lysate) for 1 hr at 4°C with mixing. Beads were washed in RIPA buffer twice, 1 M KCl, 0.1 M Na_2_CO_3_, 2 M urea (with 10 mM Tris pH 8), and twice more in RIPA buffer lacking all detergents. Beads were mixed after the last wash and bound proteins were eluted by boiling with 50 μl of 2.4x LDS loading buffer, 100 mM DTT, 15 mM biotin, 3% DMSO in three steps of 15 min each to retrieve maximal amount of biotinylated proteins. Lysates were mixed with LDS sample buffer containing 100 mm DTT and boiled for 3 min at 96°C. Mixed protein eluates were separated by one-dimensional gel electrophoresis on a 4–12% NuPage Novex Bis–Tris gel (Invitrogen), stained with the Colloidal Blue Staining Kit (Invitrogen) and digested with trypsin using in-gel digestion procedures (DOI: 10.1038/nprot.2006.468).

### SILAC-MS and data analysis

Tryptic peptides were analyzed using an EASY-nLC 1000 coupled to a Q ExactiveTM Hybrid Quadrupole-Orbitrap (Thermo Fisher Scientific). The peptides were resolved and separated on a 50 cm analytical EASY-Spray column equipped with pre-column over a 120 min gradient ranging from 8 to 38% of 0.1% formic acid in 95% acetonitrile/water. Survey full scan MS spectra (m/z 310–2000) were acquired with a resolution of 70k, an AGC target of 3 × 106 and a maximum injection time of 10 ms. Top twenty most intense peptide ions in each survey scan were sequentially isolated to an ACG target value of 5e4 with resolution of 17,500 and fragmented using normalized collision energy of 25. A dynamic exclusion of 10s and isolation width of 2 m/z were applied. SILAC peptide and protein quantification was performed with MaxQuant version 1.5.0.30 using default settings. Database searches of MS data were performed using UniProt *Canis lupus familiaris* fasta (2017_06_Dog.fasta, 25536 proteins) with tryptic specificity allowing maximum two missed cleavages, two labeled amino acids, and an initial mass tolerance of 4.5 ppm for precursor ions and 0.5 Da for fragment ions. Cysteine carbamidomethylation was searched as a fixed modification, and N-acetylation and oxidized methionine were searched as variable modifications. Labeled arginine and lysine were specified as fixed modifications and biotinylation as variable modification, depending on the prior knowledge about the parent ion. Maximum false discovery rates were set to 0.01 for both protein and peptide. Proteins were considered identified when supported by at least one unique peptide with a minimum length of seven amino acids.

### Analysis of SILAC proteomes

To generate high-confidence proximity proteomes, low quality data was filtered out; proteins with a “Ratio H/L count” < 2 and/or a “Peptide count” < 2 were not considered. All proteins tagged as “Potential contaminants”, “Reverse”, or “Only identified by site” were also removed from the analysis. In addition, non-specific streptavidin bead binders and endogenously biotinylated proteins were subtracted from all proximity proteomes. Such proteins were identified by LC-MS/MS in a control experiment in which H_2_O_2_ was omitted. The protein list was annotated with its GOCC annotations by retrieving the matching human versions from UniProt. The matching of human proteins to canine proteins was done on the gene names and verified by comparison of the protein sequences. Means of SILAC ratios were calculated by averaging the base 2 logarithmic transforms of individual experiments. Average protein intensities were calculated by dividing the peptide intensity by the number of peptides identified of the corresponding protein. Means of the average protein intensities were calculated by the averaging of the base 10 logarithmic transforms of individual experiments. Standard error of the means (s.e.m.) of both SILAC ratio and average peptide intensities were also calculated on the logarithmic transforms of individual experiments.

### Hierarchical categorization of proteomic data

Gene Ontology Cellular Component terms were grouped into broad categories representing various subcellular locations. Each protein was then grouped into a single category based on their GOCC annotations, with priority given to categories higher in a defined hierarchy. The hierarchy used was as follows: TJ, AJ, Other Junctions, Cortical & Plasma Membrane, Clathrin & Endosomal, Ciliary & Centrosomal, Cytoskeletal & Motor, Golgi & ER, Ribosomal, Mitochondrial, Cytosolic, Nuclear, Other membranes, Extracellular. Proteins with no GOCC annotations were assigned to the “Others” category. The reviewed human proteome from UniProt (release 2019_04; dated 8 May 2019) was subjected to the same hierarchical categorization in order to calculate category enrichments.

### Retrieval of protein-protein interaction data

Protein-protein interaction data was retrieved from BioGRID (version 3.5.169; dated 25 Jan 2019). Only physical protein interactions annotated in Human, Mouse, and Dog were considered. Interaction data from Affinity Capture-RNA, Protein-RNA, and Proximity Label-MS experimental methods were disregarded. BioGRID protein identifiers were converted to gene names via their UniProt IDs.

ss

### APEX2 EM sample preparation

MDCK-II cells grown on Transwell filters for 12-14 days were fixed in 0.1 M cacodylate buffer pH 7.4 (CB), 2% glutaraldehyde (GA), 2mM CaCl_2_ at room temperature for 5 min and incubated on ice in fixative for 1 hr. The diaminobenzidine (DAB) reaction was initiated by addition of freshly-diluted and filtered DAB solution (CB, 0.54 mg/ml DAB, 500 μM H_2_O_2_) and stopped after 10 min by washing five times for 2 min each in CB. Cells were post-fixed in osmium tetroxide solution (CB, 1-2% OsO_4_, 2 mM CaCl_2_) with or without counterstaining (0.5-1% K_3_[Fe(CN)_6_]), and washed five times for 2 min each in ultrapure water. Transwell filters were excised with a sharp razor blade after the osmium post-fixation washes. Cells were dehydrated in a cold graded ethanol series (20%, 50%, 70%, 90%, 100%, 100% ethanol (EMS) for 3 min each) followed by twice in 100% ethanol for 3 min each. Cells were infiltrated with a Durcupan resin:ethanol gradient (1:3, 1:1, 3:1) for 45 min each followed by 100% resin overnight. Four to six more resin changes were performed before resin-embedded samples were polymerized at 60°C for 48 hrs. Samples were sectioned into 80 nm-thick sections and mounted onto formvar- and carbon-coated slot grids. Transmission electron microscopy (TEM) was performed using a Tecnai T12 (FEI) at 120 kV. Electron micrographs were captured using a 4k×4k Eagle (FEI) CCD Camera. At least three independent experiments were performed for each cell line and images were captured randomly in at least 10 sections per sample. To produce heat-maps of APEX2 localisation, electron micrographs of identical samples recorded at the same magnification were overlaid and manually aligned in Photoshop CS6. Flattened image stacks were exported to FIJI and false-coloured.

### siRNA transfection

siRNA duplexes targeting canine Par3 or Pals1 were purchased from Dharmacon (see table 1). siRNAs were transfected at a final concentration of 100 nM using Lipofectamine RNAiMax according to the manufacturer’s recommendations (LifeTechnology). MDCK-II cells (40,000) were plated into 24 well plates in full growth medium without antibiotics and transfected the following day. Lysates for immunoblotting were prepared two-three days post-transfection (dpt). For immunofluorescence staining, transfected cells were trypsinized 2dpt and 80,000 cells were re-seeded onto filter inserts. Cells were fixed and stained by indirect immunofluorescence two days later.

### Statistical analysis of Par3 and Pals1 proximity proteomes

The Par3 and Pals1 proximity proteomes were generated in a SILAC-MS context using heavy-labelled Par3-APEX or Pals1-APEX in comparison to cytosolic APEX (NES-A2E). The proximity proteomes were thresholded by significance B testing [96] of each replicate and considering only those proteins that pass the threshold (p<0.05) in all replicates. Significance testing involves firstly characterising the distribution of normalized SILAC ratios in each replicate by the median (*r*_0_; 50 percentile) and the left (*r*_-1_; 15.87 percentile) and right (*r*_1_; 84.13 percentile) robust standard deviations. From these, the standard score of each protein (*z*) having a SILAC ratio larger than the median is calculated as:

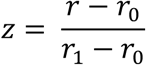

Subsequently, the z-score is used in the error function to calculate the significance

(*p*) as follows:

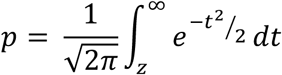

Significance B testing extends this described significance testing to account for the dependence of the standard deviation of the SILAC ratios on protein abundance. The proteins were first binned into groups of at least 300 based on their average peptide intensities. *z*-scores and *p*-values were then calculated using the *r*_0_, *r*_1_, *r*_-1_ of the subsets of proteins.

## Supplemental Figures

**Figure S1 (related to Figure 1):**
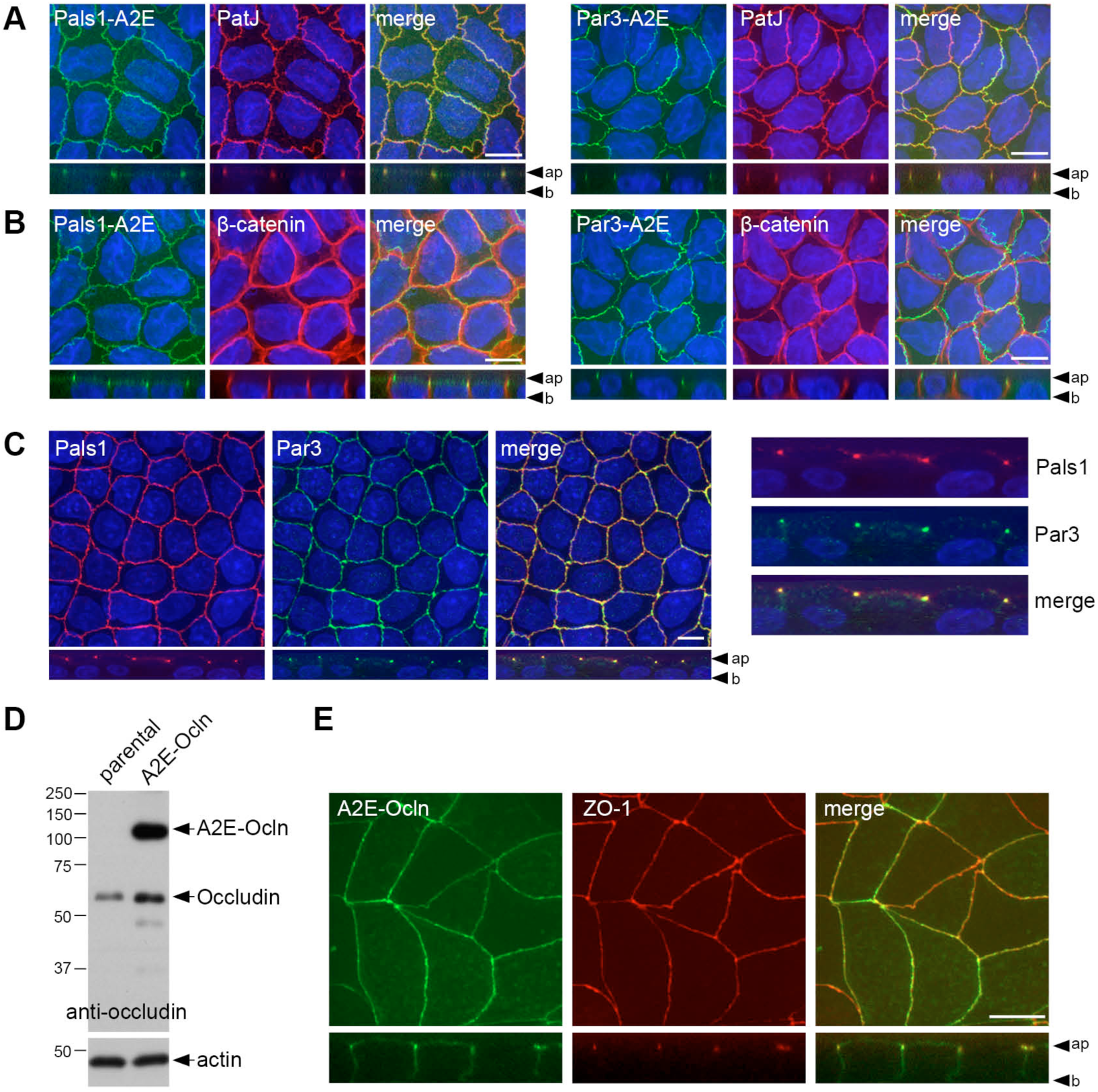
Characterization of MDCK-II cell lines stably transfected with APEX2-EGFP fusion proteins of Par3, Pals1, or occludin. **A and B.** Confocal micrographs of stable Par3-A2E and Pals1-A2E (#2) MDCK-II cells grown on filter membranes for 10 days. Cells were stained with antibodies against PatJ (A) or β-catenin (B) and counterstained with DAPI. **C.** Confocal micrographs of parental MDCK-II cells grown on filter membranes for 10 days and stained with the indicated antibodies and DAPI. **D.** Immunoblot showing the expression level of EGFP-APEX2-Occludin (A2E-Ocln) relative to endogenous occludin. Membranes were probed with the indicated antibodies. **E.** Confocal micrographs of stable A2E-Ocln cells grown on filter membranes for 10 days. Cells were stained with the indicated antibodies and DAPI. All x/y confocal micrographs are maximum intensity projections. Ap: apical; b: basal. Scale bars are 10 µm.

**Figure S2 (related to Figure 1):**
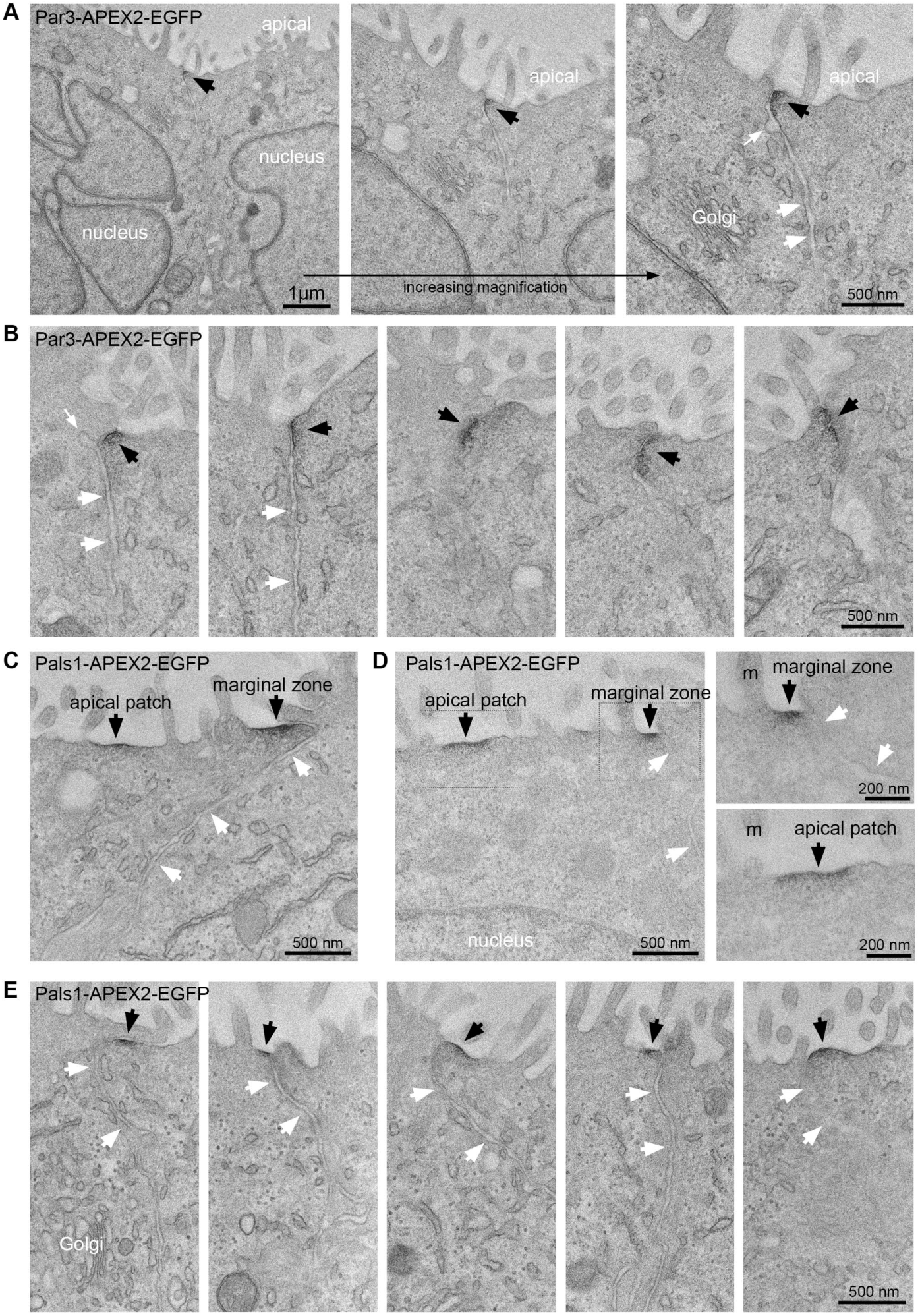
Localization of the Par3 and Pals1 APEX2-EGFP fusion proteins in counter-stained MDCK-II cells. Representative TEM micrographs of MDCK-II cells stably expressing Par3-APEX2-EGFP (A, B) or Pals1-APEX2-EGFP (C-E). Cells were counterstained with K_3_[Fe(CN)_6_] (except in D) to enhance general membrane contrast. Black arrows point to EM contrast generated by APEX2. White arrows indicate the lateral membrane. Potential endocytic structures close to the AJC are indicated with small white arrows in A and B. m=microvilli. Scale bars are indicated.

**Figure S3 (related to Figure 1):**
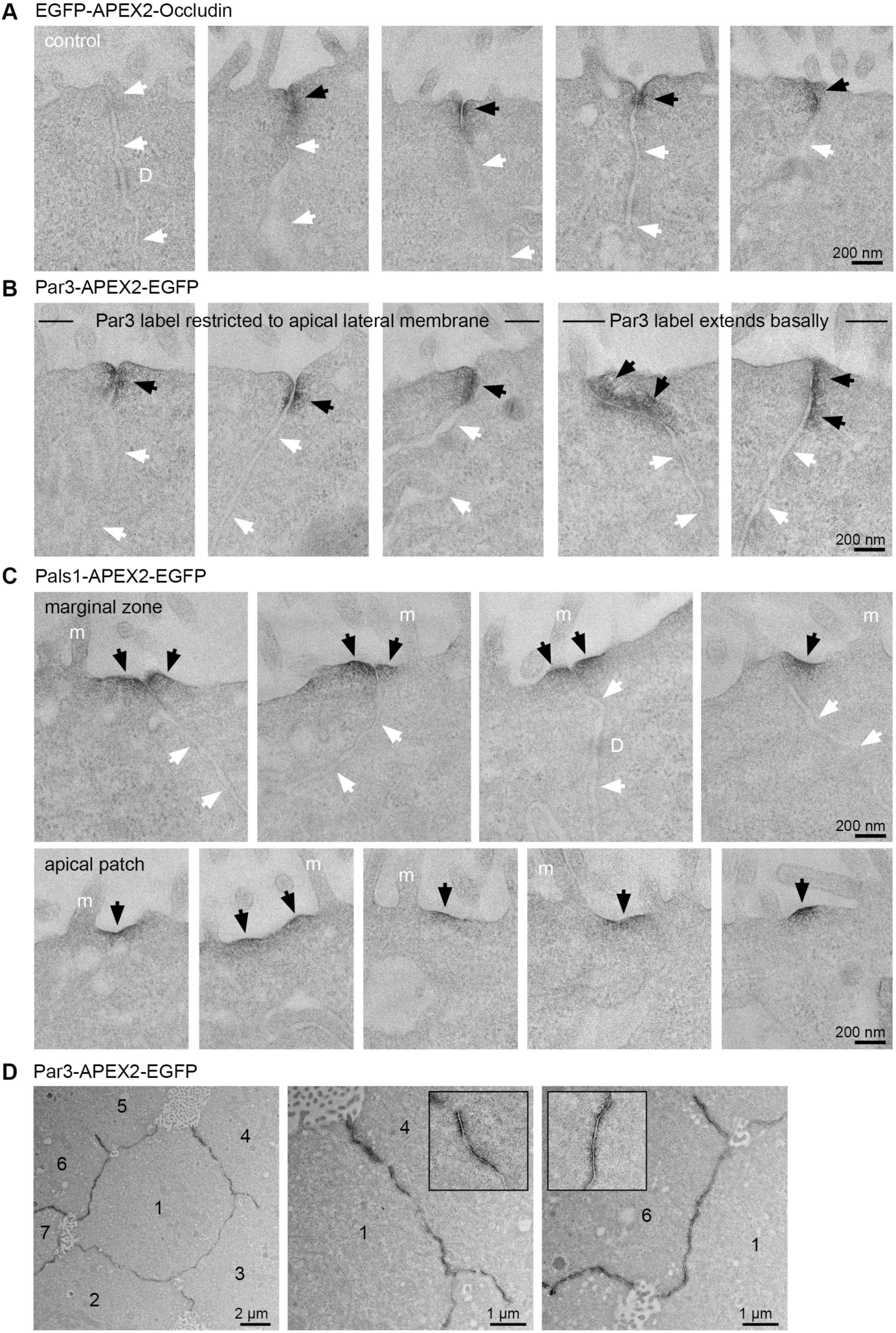
Localization of the Par3 and Pals1 APEX2-EGFP fusion proteins in unstained MDCK-II cells. Representative TEM micrographs of MDCK-II cells stably expressing EGFP-APEX2-Occludin (A), Par3-APEX2-EGFP (B) or Pals1-APEX2-EGFP (C). Cells were not counterstained. Black arrows point to EM contrast generated by APEX2. White arrows indicate the lateral membrane. D. TEM of longitudinal (*en face*) sections of Par3-A2E cells. Note the specific membrane contrast generated by Par3-A2E. m=microvilli; D=desmosome. Scale bars are indicated.

**Figure S4 (related to Figure 3):**
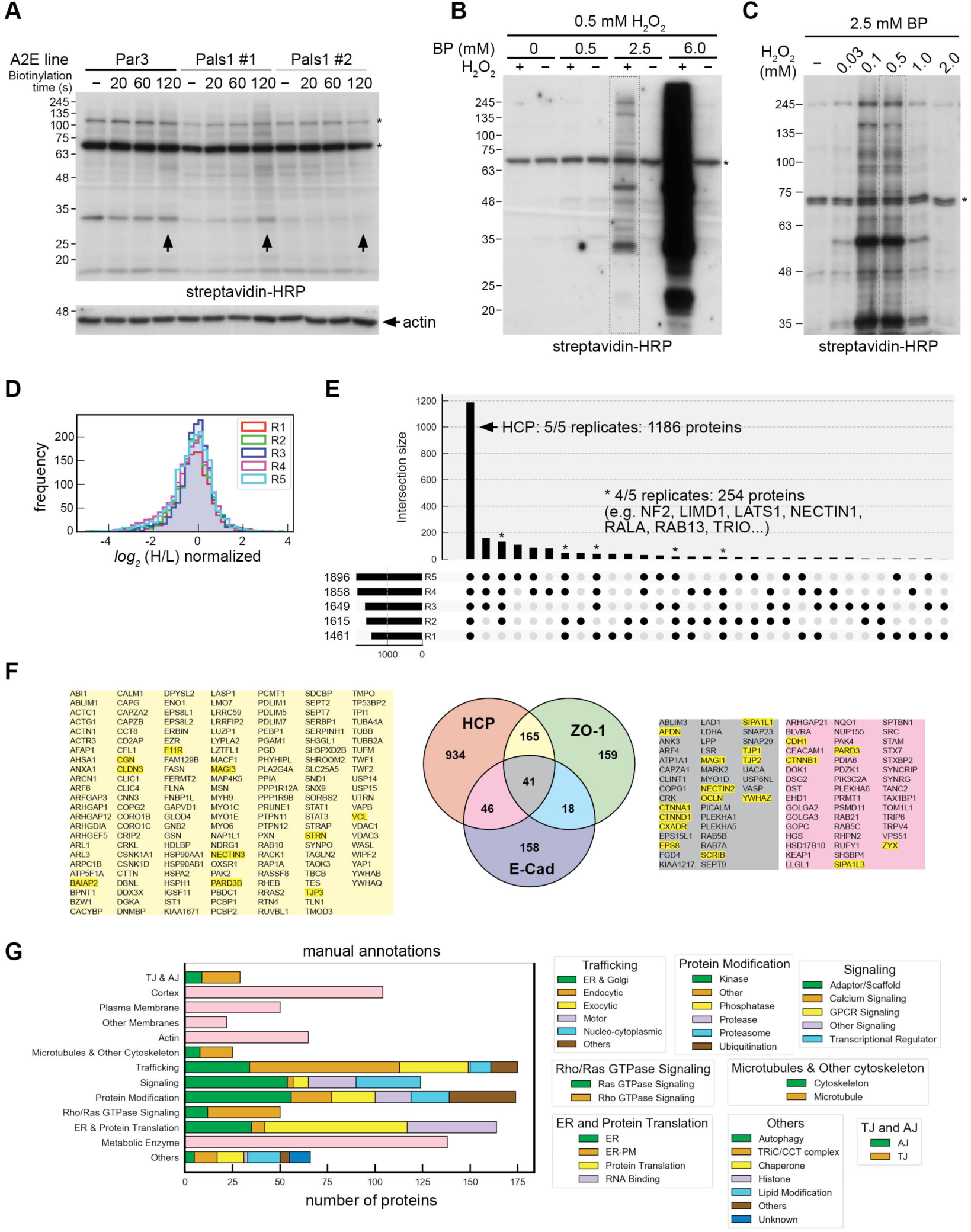
Proximity proteomics of a Par3-Pals1 APEX2 SILAC pair **A.** Streptavidin-HRP blot of cell lysates of the indicated MDCK-II A2E cell lines after 24h pre-incubation with 0.5 mM biotin phenol. The labeling reaction was initiated by incubation of cells with 1 mM H_2_O_2_ for the indicated times. Note that even after 2 min labeling the amount of biotinylated proteins observed in the respective lysates was barely above background levels (arrows). **B.** Titration of biotin phenol. Par3-A2E cells were pre-incubated for 30 min with the indicated concentrations of biotin phenol, followed by the addition of 0.5 mM H_2_O_2_ for 1 min. Cell lysates were probed with Streptavidin-HRP. **C.** Titration of H_2_O_2_. Par3-A2E cells were pre-incubated for 30 min with 2.5 mM biotin phenol, followed by the addition of H_2_O_2_ at the indicated concentrations for 1 min. Cell lysates were probed with Streptavidin-HRP. **D.** Histogram showing the distribution of the normalized *log_2_*(H/L) ratios of the identified proteins in five biological replicates (R1-R5). **E.** Histogram summarizing the frequency of protein identifications in the five replicates. 1186 proteins were identified in all five replicates and considered for a high-confidence proteome (HCP). Note that an additional 254 proteins were identified in four out of five replicates. This includes certain proteins that are expected (or known) to be localized to the AJC. **F.** Venn diagram showing common and unique proteins identified in the HCP compared to previously published ZO-1 and E-Cadherin proteomes [23, 25]. Bona-fide TJ and AJ proteins are highlighted. **G.** Manual annotation of the HCP (see Materials and Methods).

**Figure S5 (related to Figure 4):**
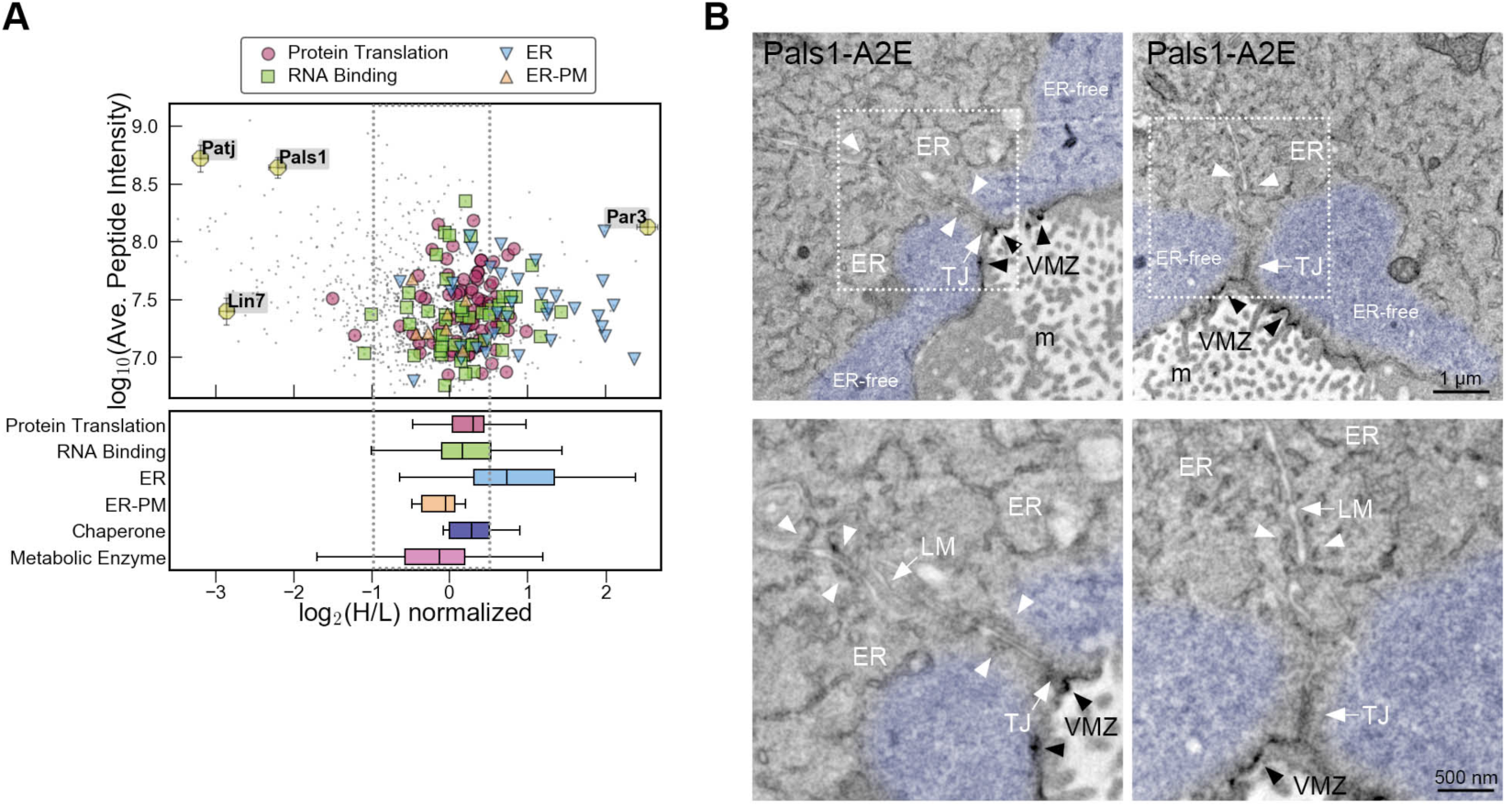
The ER is spatially associated with the basal aspect of the AJC. **A.** Scatter plot showing the apico-basal distribution of ribosomal (protein translation), RNA-binding, ER, and ER-PM contact proteins. The box plot shows the median distribution of the indicated categories including chaperones and metabolic enzymes. Note that ER proteins exhibit a basal bias. **B.** TEM micrographs of Pals1-A2E MDCK-II cells grown on filter membranes. Longitudinal (*en face*) views of the apical compartment are shown. In such sections, Pals1-A2E staining is evident as discrete membrane patches at the apical margins (black arrowheads). ER membranes (white arrowheads) are found close to apical junctions, but are absent from TJ and from the VMZ stained by Pals1-A2E. Note that the apical cytoplasm often appeared completely devoid of ER (pseudo-colored in light blue). LM=lateral membrane, m=microvilli. Scale bars are shown.

**Figure S6 (related to Figure 5):**
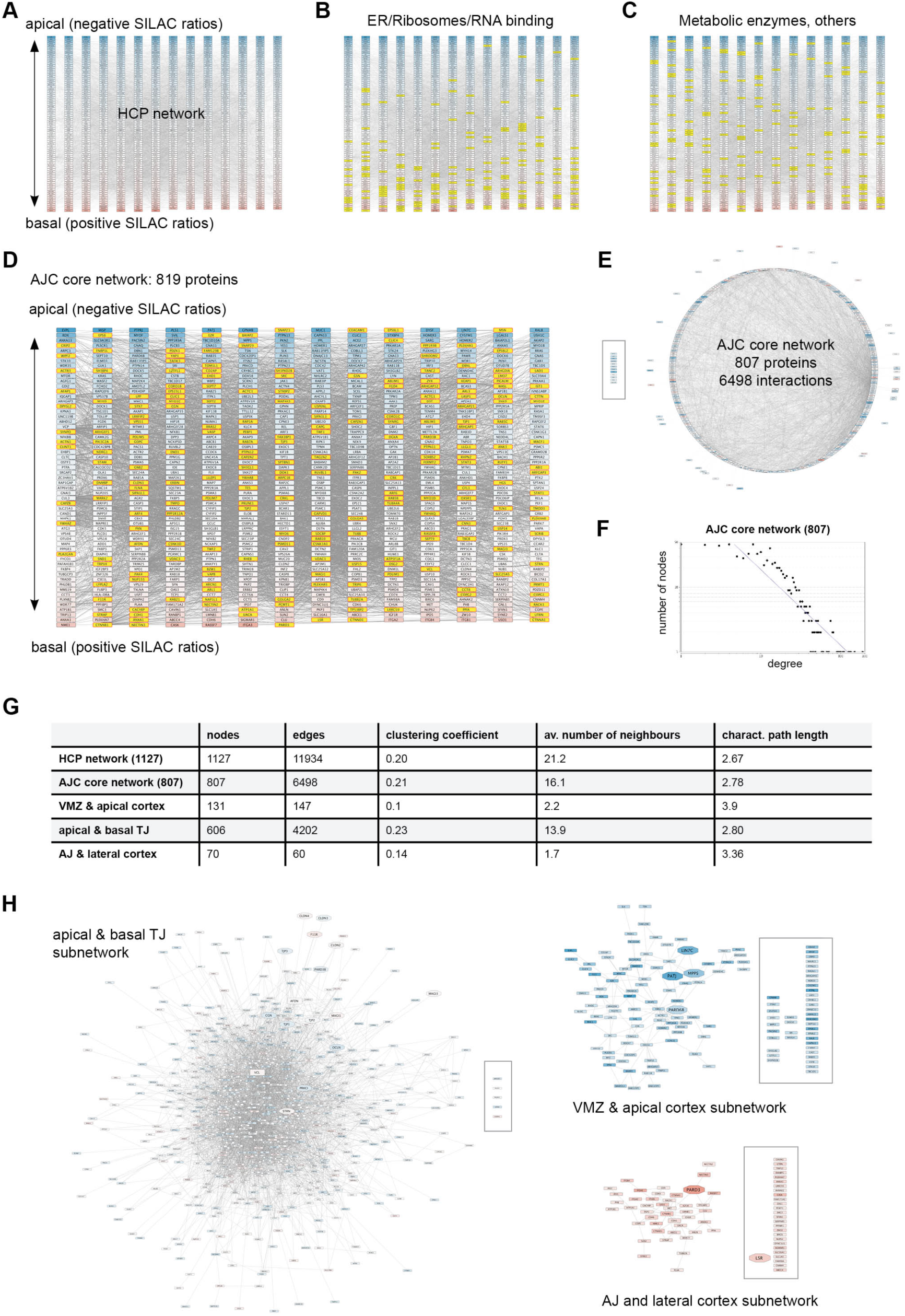
Characteristics of the AJC-associated protein interaction network. **A-C.** Cytoscape-generated group attribute layouts of the HCP interaction network (1127 proteins). PPIs were retrieved from the BioGRID database. Proteins are ranked according to their normalized *log_2_*(H/L) SILAC ratios. Proteins with negative SILAC ratios (i.e. apical) are at the top, those with positive SILAC ratios (i.e. basal) are at the bottom. **B.** ER, ribosomal and RNA binding proteins are highlighted. Note that these proteins are largely confined to the basal aspect of the network. **C.** Metabolic enzymes and other (unknown) proteins are highlighted. Note that these proteins do not show any basal or apical bias. **D.** Group attribute layout of the AJC-associated interaction network (819 proteins) after removal of proteins highlighted in B and C. Highlighted are the 220 proteins that had been identified in previous ZO-1 and E-Cadherin proteomes. **E.** Circular layout of the AJC core network shown in D. Note that only 12 proteins (boxed) did not exhibit any interactions within the network after removal of proteins shown in B and C. **F.** Node degree distribution of the AJC core network (807 proteins) determined using the network analyzer in Cytoscape. The network follows a power law of the form Y=aX^b^, with a=278.7, b=-1.172, a correlation of 0.63, and an R^2^=0.83. **G:** Table summarizing the topological characteristics of the HCP interaction network and its sub-networks determined using the network analyzer in Cytoscape. **H.** Edge-weighted spring embedded layout showing the centrality (edge betweenness) of the TJ, apical, and AJ subnetworks. Note that after elimination of apical and AJ proteins from the core AJC network, only 5 proteins (boxed) did not exhibit any interactions within the densely connected TJ subnetwork. By contrast, apical and AJ subnetworks contain a large number of proteins with no annotated interactions in these subnetworks (boxed).

**Figure S7 (related to Figure 6):**
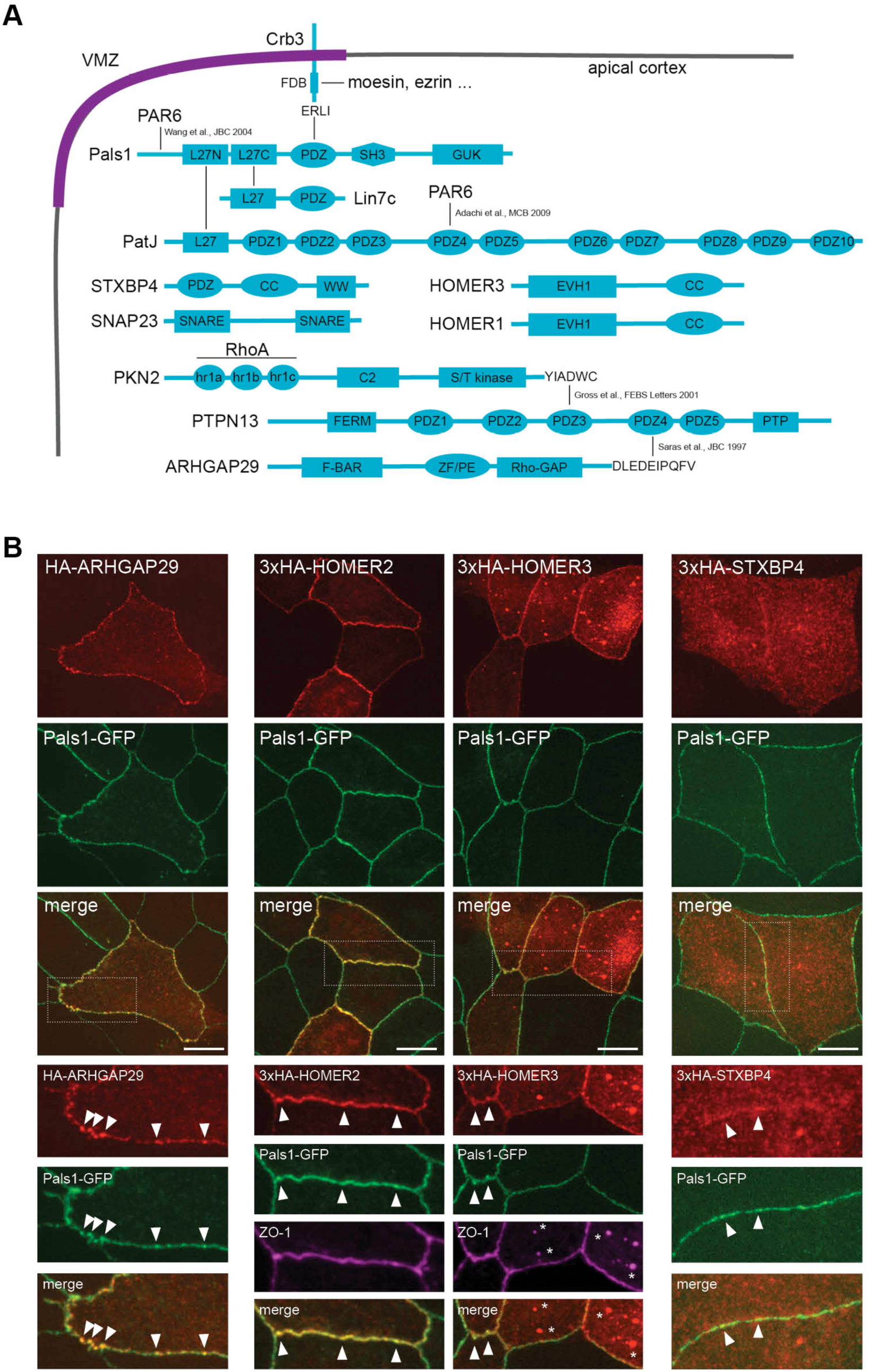
Localization of novel VMZ-associated proteins in MDCK-II cells. **A.** Domain architecture and protein-protein interactions of components of the VMZ. FDB=FERM domain binding, CC=coiled-coil domain, ZF/PE=zinc finger/ phorbol ester domain, PTP=protein tyrosine phosphatase domain, S/T kinase=serine/threonine kinase domain. **B.** Confocal micrographs of the Pals1-GFP MDCK-II rescue cell line transiently transfected with HA-tagged VMZ components. This cell line is stably transfected both with a shRNA against Pals1 and a full-length Pals1-GFP fusion protein. Defects in TJ formation observed in the parental Pals1 shRNA cell line are fully rescued by re-expression of Pals1-GFP (data not shown). Note that ARHGAP29, HOMER2, and HOMER3 colocalize with Pals1-GFP at the membrane (arrowheads). HOMER3 is also found in internal structures reminiscent of endocytic compartments (*), which contain ZO-1 but not Pals1-GFP. STXBP4 is mostly cytoplasmic and only partially localizes to the membrane. Scale bar is 10 µm.

## Supplemental Files

**Supplemental File S1:** LC-MS/MS data of the Par3-Pals1 HCP. Proteins are ranked according to their average normalized *log_2_*(H/L) ratios, with negative values (apical proteins) at the top.

**Supplemental File S2:** Manual categorization of the Par3-Pals1 HCP. Proteins are ranked according their assigned primary category.

**Supplemental File S3:** An interactive online portal of the Par3-Pals1 HCP. This html file can be opened in any internet browser. Shown is the Par3-Pals1 HCP as a scatter plot (see Figure 4A). Proteins or protein categories can be selected from the two dropdown windows. Multiple proteins and protein categories can be displayed simultaneously. The median distribution of selected protein categories is displayed as a box plot. Placing the cursor on a given protein displays the protein’s gene name, normalized *log_2_*(H/L) ratios, and average peptide intensity. Select areas of the scatter plot can be magnified using the “box zoom” tool and reset to default settings using the “reset” tool.

**Supplemental File S4:** LC-MS/MS data of the Pals1 PP. Sheet 1 shows a list of all identified proteins in the Pals1 vs NES experiment, sheet 2 shows the Pals1 PP after eliminating non-significantly enriched proteins. Proteins are ranked according to their average normalized *log_2_*(H/L) ratios, with positive values (enriched with Pals1) at the top.

**Supplemental File S5:** LC-MS/MS data of the Par3 PP. Sheet 1 shows a list of all identified proteins in the Par3 vs NES experiment, sheet 2 shows the Par3 PP after eliminating non-significantly enriched proteins. Proteins are ranked according to their average normalized *log_2_*(H/L) ratios, with positive values (enriched with Par3) at the top.

## References

1. Assemat, E., Bazellieres, E., Pallesi-Pocachard, E., Le Bivic, A., and Massey-Harroche, D. (2008). Polarity complex proteins. Biochimica et biophysica acta 1778, 614–630.

2. Rodriguez-Boulan, E., and Macara, I.G. (2014). Organization and execution of the epithelial polarity programme. Nat Rev Mol Cell Biol 15, 225–242.

3. Chen, X., and Macara, I.G. (2005). Par-3 controls tight junction assembly through the Rac exchange factor Tiam1. Nature cell biology 7, 262–269.

4. Roh, M.H., Fan, S., Liu, C.J., and Margolis, B. (2003). The Crumbs3-Pals1 complex participates in the establishment of polarity in mammalian epithelial cells. Journal of cell science 116, 2895–2906.

5. Straight, S.W., Shin, K., Fogg, V.C., Fan, S., Liu, C.J., Roh, M., and Margolis, B. (2004). Loss of PALS1 expression leads to tight junction and polarity defects. Molecular biology of the cell 15, 1981–1990.

6. Michel, D., Arsanto, J.P., Massey-Harroche, D., Beclin, C., Wijnholds, J., and Le Bivic, A. (2005). PATJ connects and stabilizes apical and lateral components of tight junctions in human intestinal cells. Journal of cell science 118, 4049–4057.

7. Shin, K., Straight, S., and Margolis, B. (2005). PATJ regulates tight junction formation and polarity in mammalian epithelial cells. The Journal of cell biology 168, 705–711.

8. Hirose, T., Izumi, Y., Nagashima, Y., Tamai-Nagai, Y., Kurihara, H., Sakai, T., Suzuki, Y., Yamanaka, T., Suzuki, A., Mizuno, K., et al. (2002). Involvement of ASIP/PAR-3 in the promotion of epithelial tight junction formation. Journal of cell science 115, 2485–2495.

9. Zihni, C., Mills, C., Matter, K., and Balda, M.S. (2016). Tight junctions: from simple barriers to multifunctional molecular gates. Nat Rev Mol Cell Biol 17, 564–580.

10. Matter, K., and Balda, M.S. (2003). Signalling to and from tight junctions. Nat Rev Mol Cell Biol 4, 225–236.

11. Tepass, U. (2012). The apical polarity protein network in Drosophila epithelial cells: regulation of polarity, junctions, morphogenesis, cell growth, and survival. Annu Rev Cell Dev Biol 28, 655–685.

12. Krahn, M.P., Buckers, J., Kastrup, L., and Wodarz, A. (2010). Formation of a Bazooka-Stardust complex is essential for plasma membrane polarity in epithelia. The Journal of cell biology 190, 751–760.

13. Benton, R., and St Johnston, D. (2003). A conserved oligomerization domain in drosophila Bazooka/PAR-3 is important for apical localization and epithelial polarity. Curr Biol 13, 1330–1334.

14. Harris, T.J., and Peifer, M. (2005). The positioning and segregation of apical cues during epithelial polarity establishment in Drosophila. The Journal of cell biology 170, 813–823.

15. Roh, M.H., Makarova, O., Liu, C.J., Shin, K., Lee, S., Laurinec, S., Goyal, M., Wiggins, R., and Margolis, B. (2002). The Maguk protein, Pals1, functions as an adapter, linking mammalian homologues of Crumbs and Discs Lost. The Journal of cell biology 157, 161–172.

16. Straight, S.W., Pieczynski, J.N., Whiteman, E.L., Liu, C.J., and Margolis, B. (2006). Mammalian lin-7 stabilizes polarity protein complexes. The Journal of biological chemistry 281, 37738–37747.

17. Lemmers, C., Medina, E., Delgrossi, M.H., Michel, D., Arsanto, J.P., and Le Bivic, A. (2002). hINADl/PATJ, a homolog of discs lost, interacts with crumbs and localizes to tight junctions in human epithelial cells. The Journal of biological chemistry 277, 25408–25415.

18. Lemmers, C., Michel, D., Lane-Guermonprez, L., Delgrossi, M.H., Medina, E., Arsanto, J.P., and Le Bivic, A. (2004). CRB3 binds directly to Par6 and regulates the morphogenesis of the tight junctions in mammalian epithelial cells. Molecular biology of the cell 15, 1324–1333.

19. Izumi, Y., Hirose, T., Tamai, Y., Hirai, S., Nagashima, Y., Fujimoto, T., Tabuse, Y., Kemphues, K.J., and Ohno, S. (1998). An atypical PKC directly associates and colocalizes at the epithelial tight junction with ASIP, a mammalian homologue of Caenorhabditis elegans polarity protein PAR-3. The Journal of cell biology 143, 95–106.

20. Zihni, C., Munro, P.M., Elbediwy, A., Keep, N.H., Terry, S.J., Harris, J., Balda, M.S., and Matter, K. (2014). Dbl3 drives Cdc42 signaling at the apical margin to regulate junction position and apical differentiation. The Journal of cell biology 204, 111–127.

21. Rees, J.S., Li, X.W., Perrett, S., Lilley, K.S., and Jackson, A.P. (2015). Protein Neighbors and Proximity Proteomics. Mol Cell Proteomics 14, 2848–2856.

22. Gingras, A.C., Abe, K.T., and Raught, B. (2018). Getting to know the neighborhood: using proximity-dependent biotinylation to characterize protein complexes and map organelles. Current opinion in chemical biology 48, 44–54.

23. Van Itallie, C.M., Aponte, A., Tietgens, A.J., Gucek, M., Fredriksson, K., and Anderson, J.M. (2013). The N and C termini of ZO-1 are surrounded by distinct proteins and functional protein networks. The Journal of biological chemistry 288, 13775–13788.

24. Fredriksson, K., Van Itallie, C.M., Aponte, A., Gucek, M., Tietgens, A.J., and Anderson, J.M. (2015). Proteomic analysis of proteins surrounding occludin and claudin-4 reveals their proximity to signaling and trafficking networks. PLoS One 10, e0117074.

25. Van Itallie, C.M., Tietgens, A.J., Aponte, A., Fredriksson, K., Fanning, A.S., Gucek, M., and Anderson, J.M. (2014). Biotin ligase tagging identifies proteins proximal to E-cadherin, including lipoma preferred partner, a regulator of epithelial cell-cell and cell-substrate adhesion. Journal of cell science 127, 885–895.

26. Guo, Z., Neilson, L.J., Zhong, H., Murray, P.S., Zanivan, S., and Zaidel-Bar, R. (2014). E-cadherin interactome complexity and robustness resolved by quantitative proteomics. Science signaling 7, rs7.

27. Pires, H.R., and Boxem, M. (2018). Mapping the Polarity Interactome. J Mol Biol 430, 3521–3544.

28. Daulat, A.M., Puvirajesinghe, T.M., Camoin, L., and Borg, J.P. (2018). Mapping Cellular Polarity Networks Using Mass Spectrometry-based Strategies. J Mol Biol 430, 3545–3564.

29. Itoh, N., Nakayama, M., Nishimura, T., Fujisue, S., Nishioka, T., Watanabe, T., and Kaibuchi, K. (2010). Identification of focal adhesion kinase (FAK) and phosphatidylinositol 3-kinase (PI3-kinase) as Par3 partners by proteomic analysis. Cytoskeleton 67, 297–308.

30. Koorman, T., Klompstra, D., van der Voet, M., Lemmens, I., Ramalho, J.J., Nieuwenhuize, S., van den Heuvel, S., Tavernier, J., Nance, J., and Boxem, M. (2016). A combined binary interaction and phenotypic map of C. elegans cell polarity proteins. Nature cell biology 18, 337–346.

31. Brajenovic, M., Joberty, G., Kuster, B., Bouwmeester, T., and Drewes, G. (2004). Comprehensive proteomic analysis of human Par protein complexes reveals an interconnected protein network. The Journal of biological chemistry 279, 12804–12811.

32. Lam, S.S., Martell, J.D., Kamer, K.J., Deerinck, T.J., Ellisman, M.H., Mootha, V.K., and Ting, A.Y. (2015). Directed evolution of APEX2 for electron microscopy and proximity labeling. Nature methods 12, 51–54.

33. Saitou, M., Ando-Akatsuka, Y., Itoh, M., Furuse, M., Inazawa, J., Fujimoto, K., and Tsukita, S. (1997). Mammalian occludin in epithelial cells: its expression and subcellular distribution. Eur J Cell Biol 73, 222–231.

34. McCarthy, K.M., Skare, I.B., Stankewich, M.C., Furuse, M., Tsukita, S., Rogers, R.A., Lynch, R.D., and Schneeberger, E.E. (1996). Occludin is a functional component of the tight junction. Journal of cell science 109 ( *Pt 9*), 2287–2298.

35. Sen, A., Sun, R., and Krahn, M.P. (2015). Localization and Function of Pals1-associated Tight Junction Protein in Drosophila Is Regulated by Two Distinct Apical Complexes. The Journal of biological chemistry 290, 13224–13233.

36. Hurd, T.W., Gao, L., Roh, M.H., Macara, I.G., and Margolis, B. (2003). Direct interaction of two polarity complexes implicated in epithelial tight junction assembly. Nature cell biology 5, 137–142.

37. Wang, Q., Hurd, T.W., and Margolis, B. (2004). Tight junction protein Par6 interacts with an evolutionarily conserved region in the amino terminus of PALS1/stardust. The Journal of biological chemistry 279, 30715–30721.

38. Adachi, M., Hamazaki, Y., Kobayashi, Y., Itoh, M., Tsukita, S., Furuse, M., and Tsukita, S. (2009). Similar and distinct properties of MUPP1 and Patj, two homologous PDZ domain-containing tight-junction proteins. Molecular and cellular biology 29, 2372–2389.

39. Hung, V., Udeshi, N.D., Lam, S.S., Loh, K.H., Cox, K.J., Pedram, K., Carr, S.A., and Ting, A.Y. (2016). Spatially resolved proteomic mapping in living cells with the engineered peroxidase APEX2. Nat Protoc 11, 456–475.

40. Shukla, P., Vogl, C., Wallner, B., Rigler, D., Muller, M., and Macho-Maschler, S. (2015). High-throughput mRNA and miRNA profiling of epithelial-mesenchymal transition in MDCK cells. BMC genomics 16, 944.

41. Citi, S., Guerrera, D., Spadaro, D., and Shah, J. (2014). Epithelial junctions and Rho family GTPases: the zonular signalosome. Small GTPases 5, 1–15.

42. Porazinski, S., Wang, H., Asaoka, Y., Behrndt, M., Miyamoto, T., Morita, H., Hata, S., Sasaki, T., Krens, S.F., Osada, Y., et al. (2015). YAP is essential for tissue tension to ensure vertebrate 3D body shape. Nature 521, 217–221.

43. Frank, S.R., Kollmann, C.P., Luong, P., Galli, G.G., Zou, L., Bernards, A., Getz, G., Calogero, R.A., Frodin, M., and Hansen, S.H. (2018). p190 RhoGAP promotes contact inhibition in epithelial cells by repressing YAP activity. The Journal of cell biology 217, 3183–3201.

44. Qiao, Y., Chen, J., Lim, Y.B., Finch-Edmondson, M.L., Seshachalam, V.P., Qin, L., Jiang, T., Low, B.C., Singh, H., Lim, C.T., et al. (2017). YAP Regulates Actin Dynamics through ARHGAP29 and Promotes Metastasis. Cell reports 19, 1495–1502.

45. Meng, Z., Qiu, Y., Lin, K.C., Kumar, A., Placone, J.K., Fang, C., Wang, K.C., Lu, S., Pan, M., Hong, A.W., et al. (2018). RAP2 mediates mechanoresponses of the Hippo pathway. Nature 560, 655–660.

46. Moya, I.M., and Halder, G. (2014). Discovering the Hippo pathway protein-protein interactome. Cell research 24, 137–138.

47. Mellacheruvu, D., Wright, Z., Couzens, A.L., Lambert, J.P., St-Denis, N.A., Li, T., Miteva, Y.V., Hauri, S., Sardiu, M.E., Low, T.Y., et al. (2013). The CRAPome: a contaminant repository for affinity purification-mass spectrometry data. Nature methods 10, 730–736.

48. Kourtidis, A., Necela, B., Lin, W.H., Lu, R., Feathers, R.W., Asmann, Y.W., Thompson, E.A., and Anastasiadis, P.Z. (2017). Cadherin complexes recruit mRNAs and RISC to regulate epithelial cell signaling. The Journal of cell biology 216, 3073–3085.

49. Huttlin, E.L., Ting, L., Bruckner, R.J., Gebreab, F., Gygi, M.P., Szpyt, J., Tam, S., Zarraga, G., Colby, G., Baltier, K., et al. (2015). The BioPlex Network: A Systematic Exploration of the Human Interactome. Cell 162, 425–440.

50. Rolland, T., Tasan, M., Charloteaux, B., Pevzner, S.J., Zhong, Q., Sahni, N., Yi, S., Lemmens, I., Fontanillo, C., Mosca, R., et al. (2014). A proteome-scale map of the human interactome network. Cell 159, 1212–1226.

51. Shannon, P., Markiel, A., Ozier, O., Baliga, N.S., Wang, J.T., Ramage, D., Amin, N., Schwikowski, B., and Ideker, T. (2003). Cytoscape: a software environment for integrated models of biomolecular interaction networks. Genome research 13, 2498–2504.

52. Assenov, Y., Ramirez, F., Schelhorn, S.E., Lengauer, T., and Albrecht, M. (2008). Computing topological parameters of biological networks. Bioinformatics 24, 282–284.

53. Zhu, X., Gerstein, M., and Snyder, M. (2007). Getting connected: analysis and principles of biological networks. Genes Dev 21, 1010–1024.

54. Reja, R., Venkatakrishnan, A.J., Lee, J., Kim, B.C., Ryu, J.W., Gong, S., Bhak, J., and Park, D. (2009). MitoInteractome: mitochondrial protein interactome database, and its application in ’aging network’ analysis. BMC genomics 10 *Suppl 3*, S20.

55. Wallace, S.W., Magalhaes, A., and Hall, A. (2011). The Rho target PRK2 regulates apical junction formation in human bronchial epithelial cells. Molecular and cellular biology 31, 81–91.

56. Fan, S., Weight, C.M., Luissint, A.C., Hilgarth, R.S., Brazil, J.C., Ettel, M., Nusrat, A., and Parkos, C.A. (2019). Role of JAM-A tyrosine phosphorylation in epithelial barrier dysfunction during intestinal inflammation. Molecular biology of the cell 30, 566–578.

57. Gross, C., Heumann, R., and Erdmann, K.S. (2001). The protein kinase C-related kinase PRK2 interacts with the protein tyrosine phosphatase PTP-BL via a novel PDZ domain binding motif. FEBS Lett 496, 101–104.

58. Saras, J., Franzen, P., Aspenstrom, P., Hellman, U., Gonez, L.J., and Heldin, C.H. (1997). A novel GTPase-activating protein for Rho interacts with a PDZ domain of the protein-tyrosine phosphatase PTPL1. The Journal of biological chemistry 272, 24333–24338.

59. Van Itallie, C.M., Lidman, K.F., Tietgens, A.J., and Anderson, J.M. (2019). Newly synthesized claudins but not occludin are added to the basal side of the tight junction. Molecular biology of the cell, mbcE19010008.

60. Staehelin, L.A. (1973). Further observations on the fine structure of freeze-cleaved tight junctions. Journal of cell science 13, 763–786.

61. Schneeberger, E.E. (1980). Heterogeneity of tight junction morphology in extrapulmonary and intrapulmonary airways of the rat. The Anatomical record 198, 193–208.

62. Van Itallie, C.M., and Anderson, J.M. (2014). Architecture of tight junctions and principles of molecular composition. Seminars in cell & developmental biology.

63. Grindstaff, K.K., Yeaman, C., Anandasabapathy, N., Hsu, S.C., Rodriguez-Boulan, E., Scheller, R.H., and Nelson, W.J. (1998). Sec6/8 complex is recruited to cell-cell contacts and specifies transport vesicle delivery to the basal-lateral membrane in epithelial cells. Cell 93, 731–740.

64. Yeaman, C., Grindstaff, K.K., and Nelson, W.J. (2004). Mechanism of recruiting Sec6/8 (exocyst) complex to the apical junctional complex during polarization of epithelial cells. Journal of cell science 117, 559–570.

65. Ahmed, S.M., and Macara, I.G. (2017). The Par3 polarity protein is an exocyst receptor essential for mammary cell survival. Nat Commun 8, 14867.

66. Nunes, F.D., Lopez, L.N., Lin, H.W., Davies, C., Azevedo, R.B., Gow, A., and Kachar, B. (2006). Distinct subdomain organization and molecular composition of a tight junction with adherens junction features. Journal of cell science 119, 4819–4827.

67. Letizia, A., Ricardo, S., Moussian, B., Martin, N., and Llimargas, M. (2013). A functional role of the extracellular domain of Crumbs in cell architecture and apicobasal polarity. Journal of cell science 126, 2157–2163.

68. Chen, C.L., Gajewski, K.M., Hamaratoglu, F., Bossuyt, W., Sansores-Garcia, L., Tao, C., and Halder, G. (2010). The apical-basal cell polarity determinant Crumbs regulates Hippo signaling in Drosophila. Proceedings of the National Academy of Sciences of the United States of America 107, 15810–15815.

69. Ling, C., Zheng, Y., Yin, F., Yu, J., Huang, J., Hong, Y., Wu, S., and Pan, D. (2010). The apical transmembrane protein Crumbs functions as a tumor suppressor that regulates Hippo signaling by binding to Expanded. Proceedings of the National Academy of Sciences of the United States of America 107, 10532–10537.

70. Robinson, B.S., Huang, J., Hong, Y., and Moberg, K.H. (2010). Crumbs regulates Salvador/Warts/Hippo signaling in Drosophila via the FERM-domain protein Expanded. Curr Biol 20, 582–590.

71. Moleirinho, S., Hoxha, S., Mandati, V., Curtale, G., Troutman, S., Ehmer, U., and Kissil, J.L. (2017). Regulation of localization and function of the transcriptional co-activator YAP by angiomotin. eLife 6.

72. Weide, T., Vollenbroker, B., Schulze, U., Djuric, I., Edeling, M., Bonse, J., Hochapfel, F., Panichkina, O., Wennmann, D.O., George, B., et al. (2017). Pals1 Haploinsufficiency Results in Proteinuria and Cyst Formation. Journal of the American Society of Nephrology : JASN 28, 2093–2107.

73. Wells, C.D., Fawcett, J.P., Traweger, A., Yamanaka, Y., Goudreault, M., Elder, K., Kulkarni, S., Gish, G., Virag, C., Lim, C., et al. (2006). A Rich1/Amot complex regulates the Cdc42 GTPase and apical-polarity proteins in epithelial cells. Cell 125, 535–548.

74. Panciera, T., Azzolin, L., Cordenonsi, M., and Piccolo, S. (2017). Mechanobiology of YAP and TAZ in physiology and disease. Nat Rev Mol Cell Biol 18, 758–770.

75. Massey-Harroche, D., Delgrossi, M.H., Lane-Guermonprez, L., Arsanto, J.P., Borg, J.P., Billaud, M., and Le Bivic, A. (2007). Evidence for a molecular link between the tuberous sclerosis complex and the Crumbs complex. Hum Mol Genet 16, 529–536.

76. Wen, J.K., Wang, Y.T., Chan, C.C., Hsieh, C.W., Liao, H.M., Hung, C.C., and Chen, G.C. (2017). Atg9 antagonizes TOR signaling to regulate intestinal cell growth and epithelial homeostasis in Drosophila. eLife 6.

77. Hansen, C.G., Ng, Y.L., Lam, W.L., Plouffe, S.W., and Guan, K.L. (2015). The Hippo pathway effectors YAP and TAZ promote cell growth by modulating amino acid signaling to mTORC1. Cell research 25, 1299–1313.

78. Santinon, G., Pocaterra, A., and Dupont, S. (2016). Control of YAP/TAZ Activity by Metabolic and Nutrient-Sensing Pathways. Trends Cell Biol 26, 289–299.

79. Zhu, M.J., Sun, X., and Du, M. (2018). AMPK in regulation of apical junctions and barrier function of intestinal epithelium. Tissue barriers 6, 1–13.

80. Wang, W., Li, X., Huang, J., Feng, L., Dolinta, K.G., and Chen, J. (2014). Defining the protein-protein interaction network of the human hippo pathway. Mol Cell Proteomics 13, 119–131.

81. Couzens, A.L., Knight, J.D., Kean, M.J., Teo, G., Weiss, A., Dunham, W.H., Lin, Z.Y., Bagshaw, R.D., Sicheri, F., Pawson, T., et al. (2013). Protein interaction network of the mammalian Hippo pathway reveals mechanisms of kinase-phosphatase interactions. Science signaling 6, rs15.

82. Post, A., Pannekoek, W.J., Ponsioen, B., Vliem, M.J., and Bos, J.L. (2015). Rap1 Spatially Controls ArhGAP29 To Inhibit Rho Signaling during Endothelial Barrier Regulation. Molecular and cellular biology 35, 2495–2502.

83. Post, A., Pannekoek, W.J., Ross, S.H., Verlaan, I., Brouwer, P.M., and Bos, J.L. (2013). Rasip1 mediates Rap1 regulation of Rho in endothelial barrier function through ArhGAP29. Proceedings of the National Academy of Sciences of the United States of America 110, 11427–11432.

84. Wu, J., Pipathsouk, A., Keizer-Gunnink, A., Fusetti, F., Alkema, W., Liu, S., Altschuler, S., Wu, L., Kortholt, A., and Weiner, O.D. (2015). Homer3 regulates the establishment of neutrophil polarity. Molecular biology of the cell 26, 1629–1639.

85. Ruby, M.A., Riedl, I., Massart, J., Ahlin, M., and Zierath, J.R. (2017). Protein kinase N2 regulates AMP kinase signaling and insulin responsiveness of glucose metabolism in skeletal muscle. American journal of physiology. Endocrinology and metabolism 313, E483–E491.

86. Yang, C.S., Melhuish, T.A., Spencer, A., Ni, L., Hao, Y., Jividen, K., Harris, T.E., Snow, C., Frierson, H.F., Jr., Wotton, D., et al. (2017). The protein kinase C super-family member PKN is regulated by mTOR and influences differentiation during prostate cancer progression. The Prostate 77, 1452–1467.

87. Galli, T., Zahraoui, A., Vaidyanathan, V.V., Raposo, G., Tian, J.M., Karin, M., Niemann, H., and Louvard, D. (1998). A novel tetanus neurotoxin-insensitive vesicle-associated membrane protein in SNARE complexes of the apical plasma membrane of epithelial cells. Molecular biology of the cell 9, 1437–1448.

88. Low, S.H., Chapin, S.J., Wimmer, C., Whiteheart, S.W., Komuves, L.G., Mostov, K.E., and Weimbs, T. (1998). The SNARE machinery is involved in apical plasma membrane trafficking in MDCK cells. The Journal of cell biology 141, 1503–1513.

89. Yu, H., Rathore, S.S., and Shen, J. (2013). Synip arrests soluble N-ethylmaleimide-sensitive factor attachment protein receptor (SNARE)-dependent membrane fusion as a selective target membrane SNARE-binding inhibitor. The Journal of biological chemistry 288, 18885–18893.

90. Wang, Q., Chen, X.W., and Margolis, B. (2007). PALS1 regulates E-cadherin trafficking in mammalian epithelial cells. Molecular biology of the cell 18, 874–885.

91. Ozcelik, M., Cotter, L., Jacob, C., Pereira, J.A., Relvas, J.B., Suter, U., and Tricaud, N. (2010). Pals1 is a major regulator of the epithelial-like polarization and the extension of the myelin sheath in peripheral nerves. J Neurosci 30, 4120–4131.

92. Fink, R.D., and Cooper, M.S. (1996). Apical membrane turnover is accelerated near cell-cell contacts in an embryonic epithelium. Dev Biol 174, 180–189.

93. Louvard, D. (1980). Apical membrane aminopeptidase appears at site of cell-cell contact in cultured kidney epithelial cells. Proceedings of the National Academy of Sciences of the United States of America 77, 4132–4136.

94. Wang, G., and Galli, T. (2018). Reciprocal link between cell biomechanics and exocytosis. Traffic 19, 741–749.

95. Roman-Fernandez, A., and Bryant, D.M. (2016). Complex Polarity: Building Multicellular Tissues Through Apical Membrane Traffic. Traffic 17, 1244–1261.

96. Cox, J., and Mann, M. (2008). MaxQuant enables high peptide identification rates, individualized p.p.b.-range mass accuracies and proteome-wide protein quantification. Nature biotechnology 26, 1367–1372.

